# Pangenomic analysis reveals plant NAD^+^ manipulation as an important virulence activity of bacterial pathogen effectors

**DOI:** 10.1101/2022.06.07.495176

**Authors:** Michelle T. Hulin, Lionel Hill, Jonathan D. G. Jones, Wenbo Ma

## Abstract

Nicotinamide adenine dinucleotide (NAD^+^) has emerged as a key component in prokaryotic and eukaryotic immune systems and the recent discovery that Toll/interleukin-1 receptor (TIR) proteins function as NAD^+^ hydrolases (NADase) link NAD^+^-derived small molecules with immune signalling. We investigated pathogen manipulation of host NAD^+^ metabolism as a virulence strategy. Using the pangenome of the model bacterial pathogen *Pseudomonas syringae,* we conducted a structure-based similarity search from 35,000 orthogroups for type III effectors (T3Es) with potential NADase activity. Thirteen T3Es, including five newly identified candidates, were identified that possess domain(s) characteristic of seven NAD^+^-hydrolyzing enzyme families. Most *P. syringae* strains that depend on the Type III secretion system to cause disease, encode at least one NAD^+^-manipulating T3E, and many have several. We experimentally confirmed the type III-dependent secretion of a novel T3E, named HopBY, which shows structural similarity to both TIR and adenosine diphosphate ribose (ADPR) cyclase. Homologs of HopBY were predicted to be type VI effectors in diverse bacterial species, indicating potential recruitment of this activity by microbial proteins secreted during various interspecies interactions. HopBY efficiently hydrolyzes NAD^+^ and specifically produces 2’cADPR, which can also be produced by TIR immune receptors of plants and other bacteria. Intriguingly, this effector promoted bacterial virulence, indicating that 2’cADPR may not be the signalling molecule that directly initiates immunity. This study highlights a host-pathogen battle ground centred around NAD^+^ metabolism and provides insight into the NAD^+^-derived molecules involved in plant immunity.

**Significance statement:** NAD^+^ metabolism plays a crucial role in plant and bacterial immunity. However, the diversity and scope of NAD^+^ processing steps in immune signalling remain unclear. Furthermore, whether pathogens can manipulate NAD^+^ metabolism to promote virulence is unknown. By conducting a pangenomic screen of the plant pathogen *P. syringae,* we found 13 type III effectors that potentially possess NADase activities, indicating that NAD^+^ manipulation is an important virulence mechanism. Further characterization of a newly identified effector HopBY showed that it produces a cyclic ADP-ribose isomer (2’cADPR) and promotes bacterial infection and symptom development. This study clarifies the role of 2’cADPR in immune signalling and provides an example of effectors as useful molecular probes to understand immunity.

## Introduction

The metabolite and redox agent nicotinamide adenine dinucleotide (NAD^+^) is a universal co-factor and redox carrier involved in diverse cellular processes (1). Recent research on the Toll/Interleukin-1 receptor (TIR) domain proteins has established a central role of NAD^+^ in immune signalling across biological kingdoms (2–4). Upon activation, TIRs cleaves NAD^+^ through a N-glycosidase activity to produce nicotinamide (NAM), ADP-ribose (ADPR) and cyclic ADPR (5). It has recently been shown that different TIRs produce distinct isomers of cADPR including 2’cADPR and 3’cADPR (6). These TIR NADase products are thought to activate immune signalling. Plant and bacterial TIRs can make some of the same cADPR isomers and plant TIRs can activate bacterial anti-viral immunity proteins *in vitro* (4). In addition to cADPR isomers, other immunity-related products of plant TIR NADase and ADP-ribosylation activities have recently been identified including pRib-AMP/ADP, ADPr-ATP and diADPR (7, 8). Therefore, NAD^+^-dependent immune signalling pathways are present in plants and bacteria (4, 9).

In plants, the TIR domain is often associated with intracellular immune receptors that also contain the nucleotide binding and leucine-rich repeat (NLR) domains (10). NLR proteins detect pathogen effectors that have entered plant cells and can promote disease. Once activated, TIR-NLR immune receptors trigger immunity through the downstream Enhanced Disease Susceptibility 1 (EDS1)-helper NLR pathways (11). Activated TIR domains have been shown to hydrolyse NAD^+^ and produce various products, including the cADPR isomer 2’cADPR (3, 6–8, 12). TIR-produced molecules pRib-AMP/ADP, ADPr-ATP and diADPR have been shown to bind to and activate specific EDS1-helper NLR complexes (7, 8), potentially linking NADase activity of activated TIR proteins to EDS1-dependent immune signalling. In addition, a TIR-only resistance protein in plants, RBA1, exhibits 2’,3’-cNMP synthase activity *in vitro* and these cyclic nucleotides may also contribute to immunity (13).

Since NAD^+^ metabolism contributes to innate immunity, we hypothesized that manipulation of this process by pathogens might promote virulence. Pathogens rely on effector proteins to manipulate host cellular processes, and effectors with NADase activities have been reported. The bacterial animal pathogen *Brucella abortus* produces TIR effectors BtpA and BtpB, which promote infection by reducing NAD^+^ levels in the animal host and BtpA has also been shown to produce 2’cADPR (5, 14). The plant pathogen *Pseudomonas syringae* produces a type III effector (T3E) HopAM1 that also has a TIR domain. HopAM1 cleaves NAD^+^ and produces 3’cADPR, which was proposed to be responsible for the immune suppression activities of HopAM1 during infection (6, 15). In addition, the bacteriophage protein Tad1 sequesters the TIR-produced molecule 2’cADPR and likely also 3’cADPR to overcome anti-viral defence of the bacterial host (6, 9, 16). In addition to producing small molecules from NAD^+^ cleavage that might interfere with host signalling pathways, the NAD^+^ reduction activities of these effectors may directly contribute to host cell death (14).

Enzymatic cleavage of NAD^+^ can occur at either the pyrophosphate bond or the N-glycosidic bond (Fig 1A). Pyrophosphate bonds can be cleaved by NUDIX (nucleoside diphosphate linked to a variable moiety X) pyrophosphatases (17) and phosphodiesterases (PDEs). The effector Avr3b produced by the oomycete pathogen *Phytophthora sojae* harbors a NUDIX domain and can hydrolyse NADH, ADPR and 2’,3’-cNMPs (13, 18). Enzymes that can cleave the N-glycosidic bond, apart from the TIR proteins, include ADP-ribosyltransferases (ART), nucleoside hydrolases (NH), ADPR cyclases, and sirtuins (Fig 1A). ARTs cleave NAD^+^ to release ADPR, which is often used for post-translational modifications of proteins and nucleic acids (19, 20). ART effectors have been identified in various pathogens although most of them are presumed to modify host protein targets (21). However, some ARTs act primarily as NAD^+^ glycohydrolases and don’t require a protein substrate (22). Conceivably, at least some ART effectors may function to manipulate NAD^+^ metabolism. The *P. syringae* T3E HopQ1 and the related *Xanthomonas euvesicatoria* T3E XopQ1 are nucleoside hydrolases. HopQ1 was shown to alter purine metabolism (23) and activate cytokinin signalling (23, 24). XopQ1 was also recently shown to hydrolyse 2’,3’-cAMP (13). So far, pathogen effectors with sirtuin or ADPR cyclase domains have not been identified.

**Figure 1.**
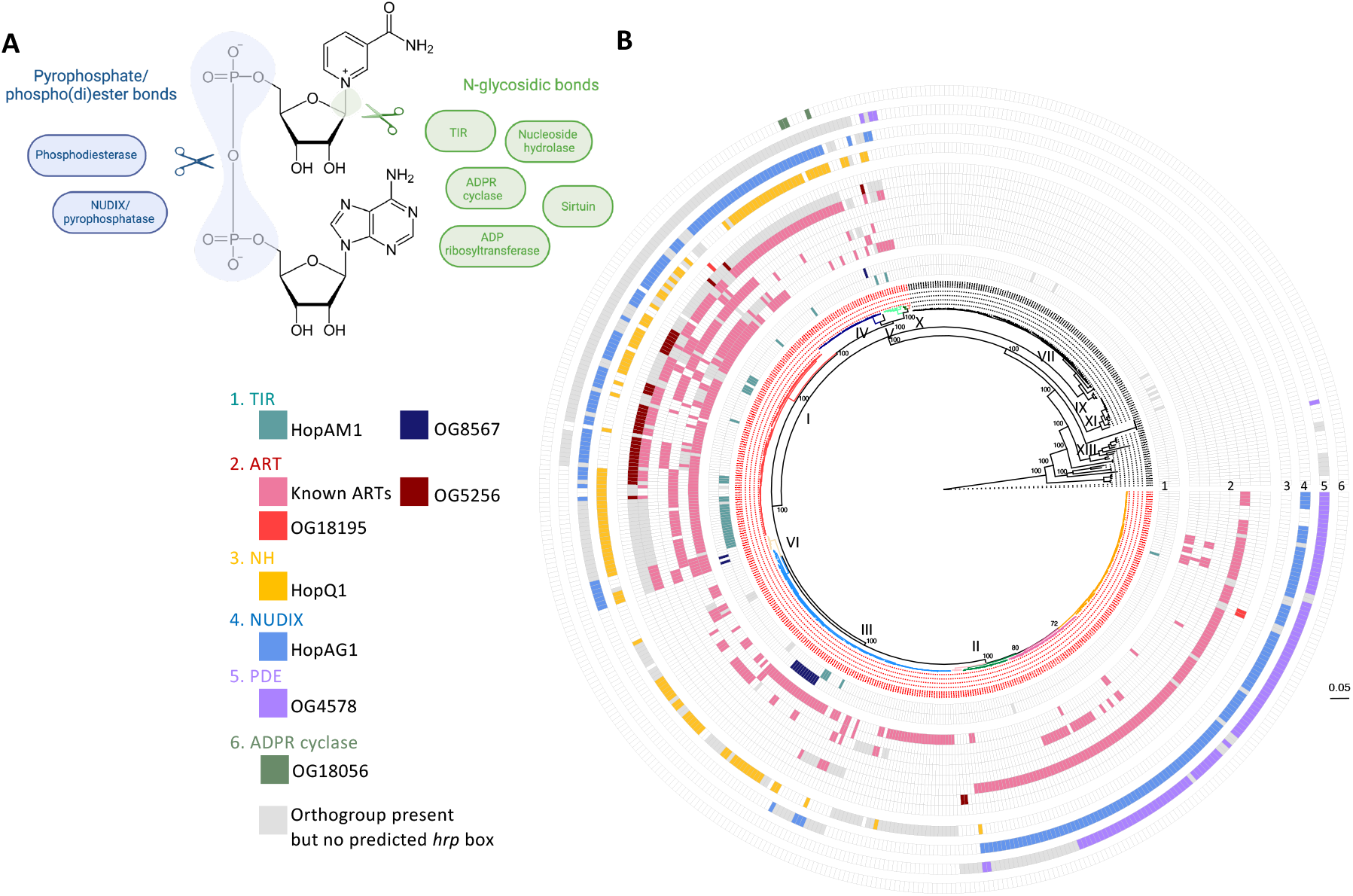
Predicted NADases as type III effectors in *P. syringae* species complex. A) Enzymes that can cleave NAD^+^. The NUDIX (nucleoside diphosphate linked to a variable moiety X) enzymes and phosphodiesterases (PDEs) cleave the pyrophosphate bond. ADP-ribosyltransferases (ART), nucleoside hydrolases (NH), ADPR cyclases, and sirtuins cleave the N-glycosidic bond. B) Predicted NADase type III effectors (T3Es) from the pangenome of *P. syringae*. Eight known T3Es and five new candidates are mapped onto the maximum-likelihood core genome phylogeny of *P. syringae* species complex. Strains in the primary phylogroups are coloured in red in the phylogenetic tree. Ring 1-6 each represents one of the six enzymatic families with NADase activity. Known T3Es with ART domains are in the order (from inside to outside) of HopF/BB, HopO1, AvrRpm, HopS, HopU1, HopAG1. Grey squares indicate the orthogroup of a specific enzymatic domain is present in the strain but does not have a putative *hrp* box in the promoter.

Here, we employ pangenomic and structure-based similarity searches to identify effectors with potential NADase activity from the species complex of the model bacterial pathogen *P. syringae*. *P. syringae* relies on T3S to delivery effectors into plant cells that suppress immunity and promote disease symptoms. We found *P. syringae* encodes T3Es with six of the seven enzymatic activities that can hydrolyze NAD^+^, suggesting that manipulation of NAD^+^ metabolism is an important virulence mechanism. We then focused on a novel T3E, named HopBY, that possesses an ADPR cyclase activity and characterized its potential virulence function. Intriguingly, HopBY hydrolyzes NAD^+^ and produces 2’cADPR *in vitro* and during infection. 2’cADPR can also be produced by plant TIR-NLRs, but our results indicate that it is unlikely to be an activator of immune signalling.

## Results

### Identification of NADases from the *P. syringae* pangenome

The *P. syringae* species complex has been divided into phylogroups, which broadly correspond to phylogenetic “species” sharing 95% Average Nucleotide Identity (ANI) (25). Plant pathogens mainly belong to the monophyletic clade of “primary” phylogroups and rely on a canonical Hrp1 type III secretion system (T3S) to cause disease (26). The “secondary” phylogroups include environment-associated strains which may not possess the Hrp1 T3S and encode very few T3Es when they do (27, 28).To examine NADase-encoding proteins across the species complex, 531 *P. syringae* genomes were analysed. A phylogeny based on the core genome grouped the strains into ≥ 95% identical ANI groups which loosely correspond to previously defined phylogroups (Fig S1). Orthology analysis assigned the pangenome of 2,867,287 protein-encoding genes into 23,583 orthogroups (OGs), with an additional 11,608 singletons. In total, 35,191 OGs were further screened for candidate NADase domains.

Two complementary approaches (HMMER and HHsearch) were employed to predict proteins that may harbour any of the seven enzymatic domains related to NAD^+^ hydrolysis (Pipeline illustrated in Fig S2A). These domains include ART, NUDIX, PDE, TIR, sirtuin, ADPR cyclase, and nucleoside hydrolase (Fig 1A; Table S1). The HMMER-based approach employed a Hidden Markov Model to identify any proteins with sequence similarity to pfam seed alignments for each enzyme group. Using this prediction method, 146 OGs were identified as candidate NADases. The HHsearch-based approach used orthogroup sequence alignments with secondary structure information. Based on HMM-HMM comparison with the PDB database, HHsearch picked up 564 OGs as candidate NADases. Importantly, 136 of the 146 candidates identified by HMMER were also identified by HHsearch (Table S2, full details in Table S3), testifying the robustness of the structure-based prediction. When mapping on the core genome phylogeny, the total of 574 candidate NADases are distributed across the *P. syringae* species complex in both primary and secondary phylogroups without a clear pattern in relation to pathogenesis (Fig S2B). This is consistent with their role in regulating housekeeping processes and anti-viral immunity.

### Prediction of NADases as Type III effectors

From the 574 candidate NADase OGs, we further identified T3Es because they are major virulence factors in *P. syringae.* Our prediction pipeline included an initial screen using the program EffectiveT3 (29) followed by a promoter search for the *hrp* box, which is bound by the T3S-specific alternative sigma factor HrpL (30). The candidates identified through this pipeline were further examined on their N-terminal amino acid sequences for characteristics related to T3S-dependent secretion (31). The outcome of this analysis revealed 13 NADase OGs as high confidence T3Es (Table 1; full list in Table S4; Fig S3-S4). The robustness of this analysis is verified by the successful identification of all the known T3Es with NADase-encoding domains including the ART effectors HopF/HopBB, HopO1, AvrRpm, HopS, and HopU1, the NUDIX effector HopAG1, the NH effector HopQ1, and the TIR effector HopAM1. The detection of multiple non-overlapping homologies to different enzymes in the same protein also revealed that HopAG1 carries a previously unknown ART domain on the N-terminus in addition to the NUDIX domain on the C-terminus. Importantly, we identified five new NADase T3E candidates, including two ARTs (OG5256, OG18195), one PDE (OG4578), one TIR (OG8567), and one ADPR cyclase (OG18056).

**Table 1.** Known NADase T3Es and predicted candidates in *P. syringae.*

| Group | Ortho-group | No. genes | No. effectors | % effectors | No. *hrp* promoter | % *hrp* promoter | No. genomes | No. T3S+ genomes | % T3S+ genomes | Notes | Secretion signal score (out of 3) |
| --- | --- | --- | --- | --- | --- | --- | --- | --- | --- | --- | --- |
| ART | 4303 | 339 | 332 | 98 | 299 | 90 | 188 | 187 | 99 | HopF/BB | 2 |
| ART | 5256 | 144 | 130 | 90 | 38 | 29 | 37 | 35 | 95 | Novel | 3 |
| ART | 5397 | 133 | 91 | 68 | 54 | 59 | 46 | 46 | 100 | HopO1 | 3 |
| ART | 5686 | 112 | 103 | 92 | 91 | 88 | 87 | 87 | 100 | AvrRpm | 3 |
| ART | 6340 | 69 | 54 | 78 | 53 | 98 | 53 | 53 | 100 | HopS | 3 |
| ART | 9885 | 14 | 14 | 100 | 10 | 71 | 10 | 10 | 100 | HopU1 | 3 |
| ART | 18195 | 3 | 3 | 100 | 3 | 100 | 3 | 3 | 100 | Novel | 3 |
| ART<sup>a</sup> | 4463 | 297 | 223 | 75 | 213 | 96 | 213 | 213 | 100 | HopAG1 | 3 |
| NH | 4871 | 195 | 124 | 64 | 121 | 98 | 118 | 117 | 99 | HopQ1 | 2 |
| NUDIX | 4463 | 297 | 223 | 75 | 213 | 96 | 213 | 213 | 100 | HopAG1 | 3 |
| PDE | 4578 | 258 | 214 | 83 | 102 | 48 | 101 | 100 | 99 | Novel | 2 |
| TIR | 6638 | 55 | 49 | 89 | 45 | 92 | 38 | 37 | 97 | HopAM1 | 3 |
| TIR | 8567 | 23 | 21 | 91 | 15 | 71 | 14 | 14 | 100 | Novel | 2 |
| ADPR cyclase | 18056 | 3 | 3 | 100 | 3 | 100 | 3 | 3 | 100 | Novel | 3 |

No. genes: number of genes in orthogroup.
No. effectors: number of genes within orthogroup that were predicted to encode an effector.
% effectors: % orthogroup that may encode effector.
No. *hrp* promoter: Number of predicted effector genes per orthogroup that had a predicted *hrp* box
% *hrp* promoter: % effector genes in orthogroup that had *hrp* promoter.
No. T3S+ genomes: Number of genomes encoding the genes in each orthogroup predicted as effectors with a *hrp* box which have the genes required for Hrp1 T3S machinery.
% genomes with Hrp1 T3S: % of effector genes with a *hrp* box per orthogroup that were in genomes encoding Hrp1 T3S.
Secretion signal score: A score from 0-3 based on three previously defined criteria of T3E secretion signals: 1) a high % serines; 2) a lack of negative amino acids in the first 12 residues; 3) an aliphatic amino acid at position 3 or 4.
<sup>a</sup> Although the main hit of OG0004463 (HopAG1) was a NUDIX domain, a further domain with homology to ART was predicted at a different region of the protein.

The distribution of the 13 known NADase T3Es and new candidates was mapped on the core genome phylogeny of *P. syringae* species complex (Fig 1B). A clear difference between the primary and secondary phylogroups can be visualized, with known and predicted NADase T3Es almost exclusively encoded by strains belonging to the primary phylogroups, consistent with their potential role in host manipulation (Fig 1B and Fig S5). 93% of the primary phylogroup strains have at least one NADase T3Es and 72% possess T3Es belonging to more than one enzyme family. On average, each primary phylogroup strain encodes three T3Es that potentially hydrolyze NAD^+^. NAD^+^ hydrolysis is therefore a common virulence mechanism utilized by *P. syringae* pathogens.

### OG18056 is a NADase with structural homology to both ADPR cyclases and TIRs

Considering the essential role of NAD-derived cADPR isomers in plant immunity, we were particularly interested in a novel T3E candidate OG18056, which is predicted to have an ADPR cyclase domain that has not been found in pathogen effectors. ADPR cyclase shares a similar enzymatic activity with TIR and can also hydrolyze NAD^+^ to produce cADPR (32). Using AlphaFold2 (33), we generated a structural model for OG18056. Superimposition of this model with the human ADPR cyclase CD38 showed an RMSD value of 2.94 Å. The structural model of OG18056 also showed similarity with the TIR domain of the human SARM1 protein with an RMSD value of 2.96 Å (Fig 2A). These results suggest that OG18056 possesses a fold that resembles both TIR and ADPR cyclase, although our initial prediction only revealed similarity with ADPR cyclase through HHsearch structure-based analysis.

**Figure 2.**
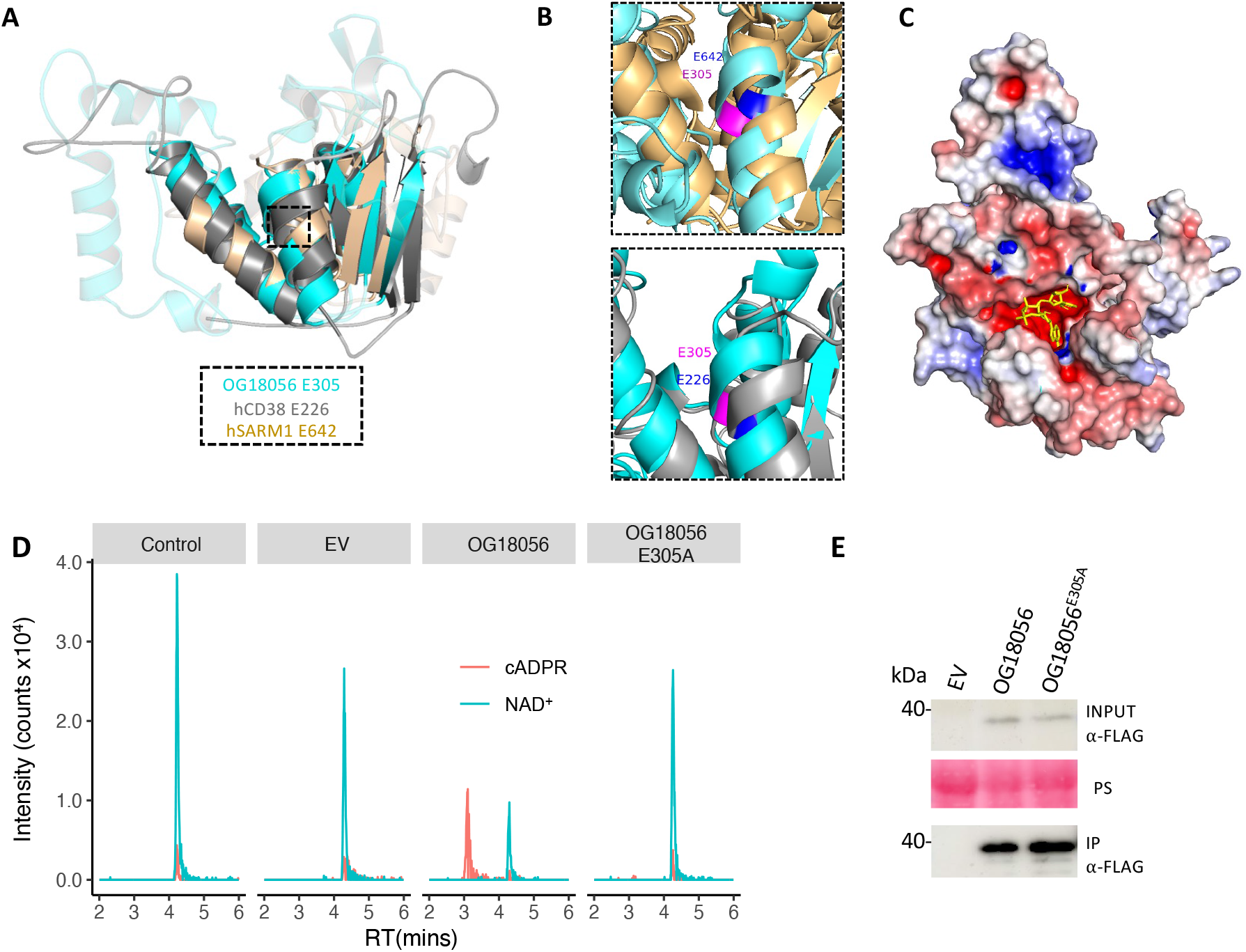
OG18056 is an active NADase that produces a variant of cADPR. A) Alignment of a structural model of OG18056 (predicted by AlphaFold2, in cyan) with a human ADPR cyclase CD38 (6VUA, in grey) and the TIR domain of human SARM1 (6O0R, 561-700 aa, in light brown). For visualisation purposes, sequences outside the alignment region were removed and those that did not align within the region are more transparent. The position of the predicted catalytic Glu residues, which are aligned in the three proteins, is highlighted with a box. B) A close-up of alignments of OG18056 with TIR-hSARM1 (upper) and hCD38 (lower) at the predicted catalytic site showing a Glu E305 in OG18056 that is aligned with the catalytic residues E642 in TIR-hSARM1 and E226 in hCD38. C) Electrostatic image of the surface of the structure model of OG18056 with regions of positive, neutral, and negative charge coloured in blue, white, and red respectively. A molecule of NAD^+^ was docked into the predicted catalytic pocket. D) HPLC chromatograms showing the cADPR product produced by OG18056 using a semi-*in vitro* assay. OG18056 and OG18056^E305A^ were expressed in *N. benthamiana.* After immuno-precipitation, the proteins were incubated with NAD^+^ for one hour before subjected to LCMS analysis. The controls were NAD^+^ only without addition of beads and immunoprecipitation of proteins from leaves infiltrated with *Agrobacterium* carrying an empty vector (EV). Peaks corresponding to m/z 542.06 (cADPR) and m/z 664.11 (NAD^+^) were identified. E) Immunoblots detecting the expression of OG18056 and OG18056^E305A^ in *N. benthamiana* (input) and after immunoprecipitated (IP). The IPed samples were used for the LCMS assays. OG18056 and OG18056^E305A^ were tagged with C-terminal 3xFLAG. Empty vector (EV) was included as a control. Ponceau Staining (PS) of the membrane was used as a loading control.

The NAD^+^ hydrolysis activity requires a catalytic glutamate (Glu, E) in TIR and ADPR cyclase. Structural alignment revealed E305 in OG18056 to be located at the corresponding position of the catalytic E642 in hSARM1 (2) as well as E226 in hCD38 (34) (Fig 2B). Furthermore, a ligand-binding pocket, containing the putative catalytic residue E305, was predicted in OG18056 3D model using the program P2RANK with a probability of 0.9 (35). Molecular docking revealed that NAD^+^ could fit into this predicted pocket with a binding affinity of −8.76 kcal/mol (Fig 2C). These results are consistent with the hypothesis that OG18056 is an active NAD^+^-cleaving enzyme.

To evaluate the enzymatic activity of OG18056, we used HPLC-MS analysis to detect NAD^+^ hydrolysis and production of various form(s) of cADPR. In a semi-*in vitro* assay (36), 3xFLAG-tagged OG18056 and its predicted catalytic mutant OG18056^E305A^ were expressed in *Nicotiana benthamiana* by *Agrobacterium-mediated* transient expression. Wild type and mutant OG18056 proteins were immunoprecipitated and incubated with NAD^+^. The resulting metabolites were extracted and analyzed by LCMS. The results show that OG18056 produced a peak for m/z of 542.06, which is consistent with that of cADPR, at a retention time of ~3.0 minutes (Fig 2D). OG18056^E305A^ did not produce this compound. The proteins were confirmed to be expressed and immunoprecipitated (Fig 2E). These results demonstrate that OG18056 is indeed an active NADase that produces a mass corresponding to cADPR.

### OG18056 (HopBY) produces 2’cADPR during bacterial infection

To assess whether OG18056 is a secreted protein dependent on T3S, we tested its expression pattern and T3S-dependent secretion using a minimal medium (Fig 3A). The gene encoding OG18056, tagged at the C-terminus with a HA epitope, was introduced into *P. syringae* pv. tomato strain DC3000 (PstDC3000) under its native promoter. Bacterial cells were induced in a *hrp*-minimal medium (HM) and OG18056 proteins were detected in the cell culture supernatant by western blotting. We observed that OG18056 had a minimal expression in the rich medium KB, but was drastically induced in the HM medium, an expression pattern commonly observed in T3S-related genes. Consistent with the detection of a *hrp* box in the promoter of OG18056 gene, this result shows that OG18056 is regulated by T3S. Furthermore, OG18056 proteins were detected in the supernatant of PstDC3000 cell culture, but not in the supernatant of the *DhrcC* mutant of PstDC3000, which does not form a functional T3S injectosome. These results suggest that OG18056 is likely a T3E that can be secreted in a T3S-dependent manner. We therefore name this new T3E HopBY based on the *P. syringae* T3E nomenclature system (28, 37).

**Figure 3:**
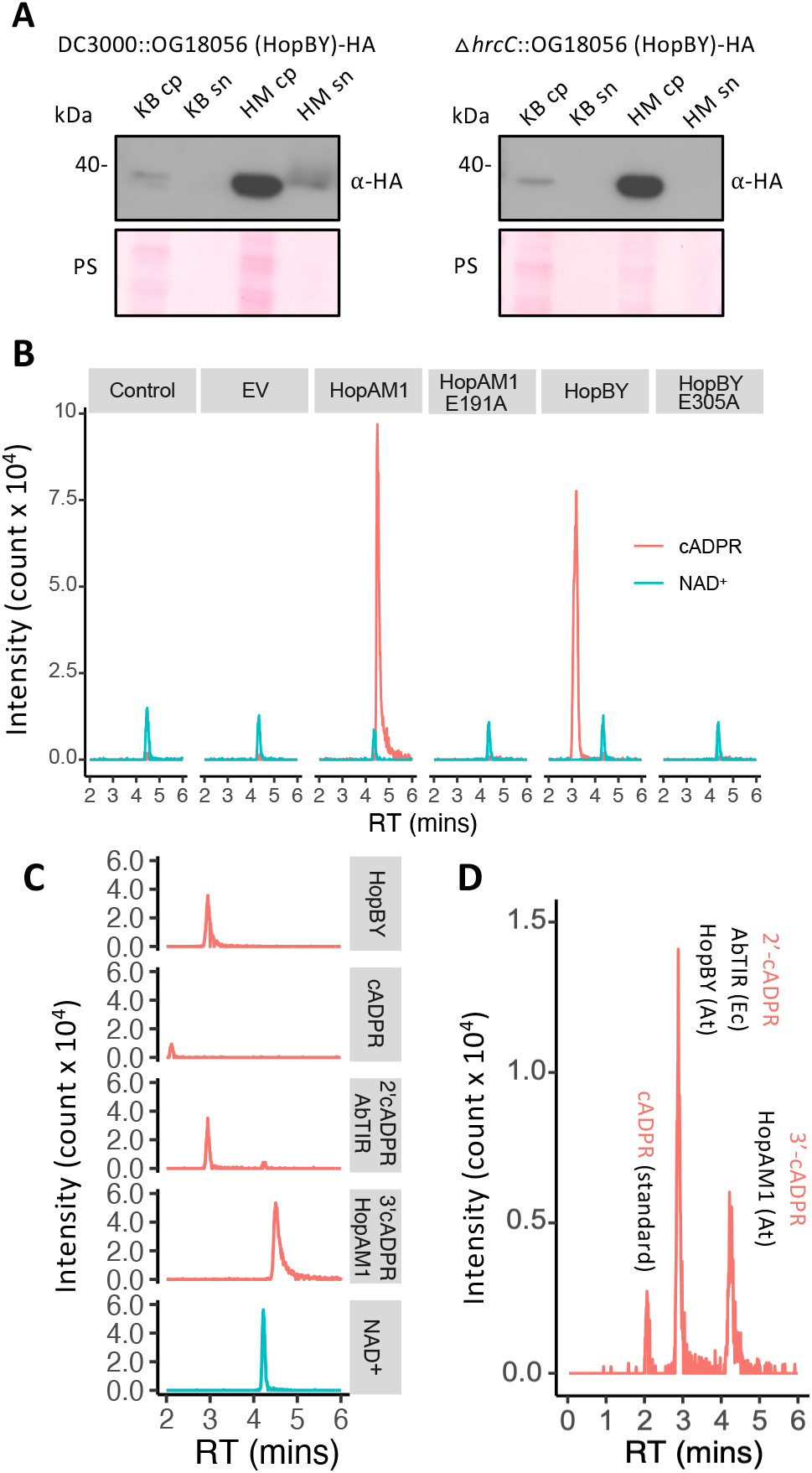
OG18056 (HopBY) is a true type III effector that produces 2’cADPR during bacterial infection. A) Immunoblots showing the induction and secretion of OG18056 in a T3S-dependent manner by *P. syringae.* OG18056-HA was introduced in PstDC3000 or the T3S mutant Δ*hrcC* under its native promoter. The bacteria were cultured in the rich medium King’s B (KB) or a Hrp-inducing minimal medium (HM). Proteins were extracted from cell pellet (cp) or supernatant (sn) and OG18056 was detected by western blotting using an anti-HA antibody. Ponceau Staining (PS) of the membrane was used as a loading control. OG18056 is hereafter referred to as HopBY. B) HPLC chromatograms of *A. thaliana* leaf extracts using tissues 18 hours post-inoculation with the *P. syringae* “effectorless” mutant D36E expressing HopBY. Leaves treated with 10mM MgCl2 was used as the control. Leaves inoculated with D36E carrying an empty vector (EV), or expressing HopBY^E305A^, HopAM1 or HopAM1^E191A^ were included for comparison. All effectors were cloned under their native promoter. Peaks corresponding to m/z 542.06 (cADPR) and m/z 664.11 (NAD^+^) were identified. C) HPLC chromatograms of different cADPR variants compared to the product of HopBY (extracted from *A. thaliana* inoculated with D36E expressing HopBY). 2’cADPR was detected from cell lysate of *E. coli* expressing AbTIR, and 3’cADPR was detected from *A. thaliana* leaves inoculated with D36E expressing HopAM1. cADPR and NAD^+^ were purchased from Sigma. D) HPLC chromatogram of a mixture of samples as described in Panel C. Only one peak appeared at a retention time 3.0 min, indicating that the HopBY product is likely 2’cADPR. At: *Arabidopsis thaliana*; Ec: *E. coli*

We next examined whether HopBY hydrolyzes NAD^+^ and produces isomer(s) of cADPR during natural infection. HopBY or HopBY^E305A^ was expressed in the D36E mutant of PstDC3000 under its native promoter (Fig S6). D36E has the intact T3S but lacks all 36 known T3Es, and is not pathogenic (38). The transformed bacteria were then used to inoculate *Arabidopsis thaliana.* Eighteen hours post-infiltration, metabolites were extracted from the inoculated leaf tissues and subjected to LCMS analysis. We detected a clear peak corresponding to m/z 542.06 at the same retention time (~3 min) as in the semi-*in vitro* assay (Fig 3B). This peak was absent in leaves inoculated with D36E carrying the empty vector or expressing the catalytic mutant HopBY^E305A^, indicating that it is the product of HopBY NADase activity. As a control, leaves inoculated with D36E expressing HopAM1 showed a peak with a retention time of ~4.5 minutes, corresponding to 3’cADPR, as previously reported (6, 15). To determine which isomer of cADPR was produced by HopBY, we ran LCMS on a cell lysate of *E. coli* expressing AbTIR, which is produced by the opportunistic bacterial pathogen *Acinetobacter baumannii.* AbTIR is known to produce 2’cADPR (6). Interestingly, the HopBY product had the same retention time as the AbTIR product (Fig 3C). When running a mixture of the different cADPR samples, a single peak with m/z of 542.06 and a retention time of ~3.0 minutes appeared, suggesting that the NAD^+^ hydrolysis product of HopBY is likely the same or very similar to AbTIR, i.e. 2’cADPR (Fig 3D). Comparison of MS2 fragmentation profiles demonstrated that the peak produced by HopBY generated similar fragment ions to the cADPR standard and cADPR variants produced by other NADases, confirming that the HopBY product was indeed a cADPR (Fig S7).

To determine if HopBY has any impact on overall NAD^+^ levels during infection, we measured the abundance of NAD^+^ at days 0 and day 3 after inoculation with D36E carrying either HopBY or HopAM1 (5 x 10^6^ CFU/mL). NAD^+^ levels showed a small but significant reduction in leaf tissues inoculated with HopBY-expressing bacterium at day 3 compared to day 0 (Fig. S8). This reduction was not observed in tissues inoculated with D36E expressing HopAM1 or carrying the empty vector. These results suggest that HopBY may reduce NAD^+^ levels in host cells as a virulence mechanism.

### HopBY promotes bacterial infection and necrosis

We next examined the potential virulence activity of HopBY during bacterial infection. D36E expressing HopBY or the catalytic mutant HopBY^E305A^ was used to inoculate *A. thaliana* plants at a dose of 5 x 10^6^ CFU/mL. We observed chlorotic symptoms in leaves inoculated with D36E expressing HopBY, but not HopBY^E305A^, after three days (Fig 4A). When inoculated at a higher dose (1 x 10^8^ CFU/mL), tissue collapse appeared at 24 hours post inoculation, which is similar in appearance to immunity-related hypersensitive response triggered by effectors recognized by NLRs, such as AvrRps4 (Fig 4B). It is noteworthy that D36E expressing AvrRps4 did not induce the chlorosis at the lower inoculum as HopBY did (Fig 4A). Similar to chlorosis, the cell death-like phenotype was dependent on the NADase activity as the HopBY^E305A^ mutant failed to induce this symptom. In addition, inoculation of *Pst* DC3000Δ*hrcC* expressing HopBY did not cause these symptoms, which is consistent with the previous results suggesting that HopBY is likely translocated during bacterial infection in a T3S-dependent manner (Fig. S9).

**Figure 4:**
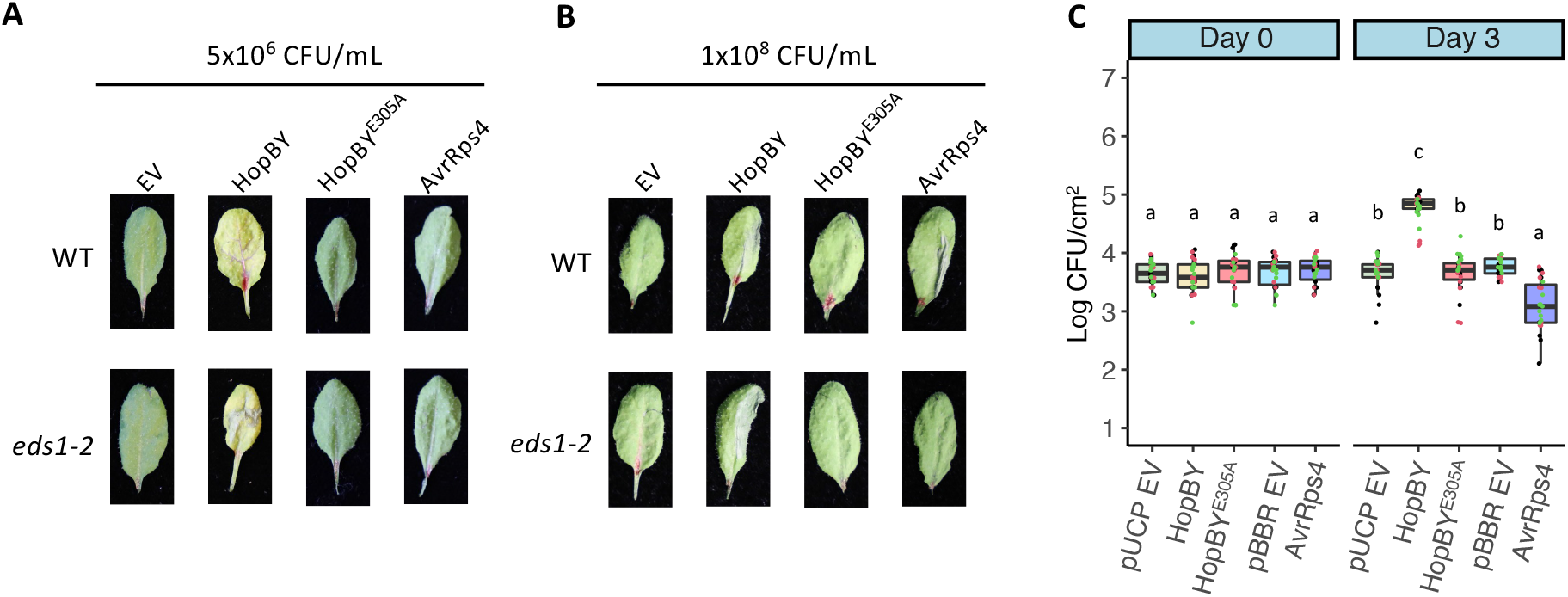
HopBY enhanced bacterial growth and triggered disease-like symptoms in *A. thaliana*. A) Disease symptom development in *A. thaliana* inoculated by the *P. syringae* mutant D36E carrying the empty vector (EV), or expressing HopBY, HopBY^E305A^ or AvrRps4. Adult leaves of wildtype (WT) and *eds1-2* mutant were inoculated with bacterial suspensions of 5 x 10^6^ CFU/mL and symptoms were imaged at 3 days post inoculation. B) Cell death-like phenotype induced by HopBY at a high inoculum dose (5 x 10^8^ CFU/mL) in an EDS1-independent manner. D36E expressing individual effectors were used to inoculate *A. thaliana* adult leaves, which were imaged one day post inoculation. C) Bacterial populations (in Log CFU/cm^2^) measured at 0 and 3 days post inoculation in wildtype *A. thaliana*. D36E transformants were used to inoculate adult leaves at 5 x10^5^ CFU/mL. Individual data points with different colors represent results from three separate experiments. Tukey-HSD significance groups extracted from ANOVA are presented (p<0.05). HopBY was cloned in the vector pUCP20tk but AvrRps4 was cloned in the vector pBBR1MCS-5. Therefore both empty vectors (pUCP EV and pBBR EV) were included as controls.

Because 2’cADPR can be produced by some plant TIR-NLRs, it is therefore a possibility that HopBY may activate an immune response, which would lead to cell death due to hypersensitive response. To examine this, we inoculated D36E expressing HopBY1 in the *A. thaliana* mutant *eds1-2,* which is critical for TIR-NLR-mediated immune signalling (39). The results show that HopBY was still able to induce the chlorosis (Fig 4A) and cell death-like phenotypes in the *eds1* mutant (Fig 4B). In contrast, the cell death induced by AvrRps4 as a hypersensitive response was abolished in *eds1-2* (Fig 4B). To further confirm HopBY did not trigger immunity in *A. thaliana*, we evaluated the *in planta* bacterial population at three days post inoculation. While expression of AvrRps4 caused a decline in bacterial population, HopBY actually promoted infection (Fig 4C). This virulence activity was not observed in the catalytic mutant HopBY^E305A^. These results demonstrate that HopBY enhances bacterial infection by promoting bacterial colonization and symptom development. These virulence functions rely on its NADase activity, and possibly related to the production of 2’cADPR.

### HopBY homologues are associated with T3 and T6 secretion systems

HopBY is only present in three *P. syringae pv. coronafaciens* strains, which are pathogens of grass species. The *hopBY* gene is flanked by transposase genes and nearby genes encoding conjugation machinery *(virB* homologues). It is within 15 Kb of T3E homologues *hopV1 and hopAA1* as well as a coronatine biosynthesis operon, which also contributes to virulence (Fig S10A). This genomic location suggests that *hopBY* may have been acquired via horizontal gene transfer. A skew in GC content further indicates that this genomic region may represent a pathogenicity island (Fig S10A).

Using BLASTP, we identified homologs of HopBY in diverse bacterial lineages ranging in amino acid sequence identity from 26%-100%. HHsearch confirmed that these homologs all have an ADPR cyclase-like fold on their C terminus (Table S5). Examination of the genomic region surrounding all homologs revealed they are often associated with transposons, indicating a strong signature of horizontal gene transfer (Fig S10B). A phylogeny based on structure-guided protein sequence alignment grouped these HopBY homologs into two clades (Fig 5A). The HopBY from *P. syringae* is grouped with homologs from other *Pseudomonas* spp. as well as plant pathogenic Enterobacteriaceae species within the genera *Brenneria* and *Pectobacterium.* All these genomes contain genes encoding a Hrp1 T3S, the canonical T3S found in *P. syringae.* Importantly, the homologous genes in this clade are all preceded by a *hrp-* box promoter (Table S5), indicating that they are likely T3Es. In the *Brenneria* (n=2) genomes, the *hopBY* homologous genes are present close to the T3S machinery gene clusters, further supporting their identity as T3Es. The other clade includes more distinct HopBY homologs including one in *Paraburkholderia* sp. ZP32-5, which was also predicted as a possible T3E. *Paraburkholderia* sp. ZP32-5 contains the Hrp2 type of T3S machinery and the HopBY homolog gene is located in close proximity to the T3S gene cluster. Upstream of this gene there are PIP-box motifs, which are associated with Hrp2 T3S-regulated gene expression. Therefore, this HopBY homolog is likely also a T3E. Interestingly, other homologs in this clade are often located adjacent to type VI secretion system (T6S) genes such as *vgrG, tssA, vasA* and *tssG.*

**Fig 5:**
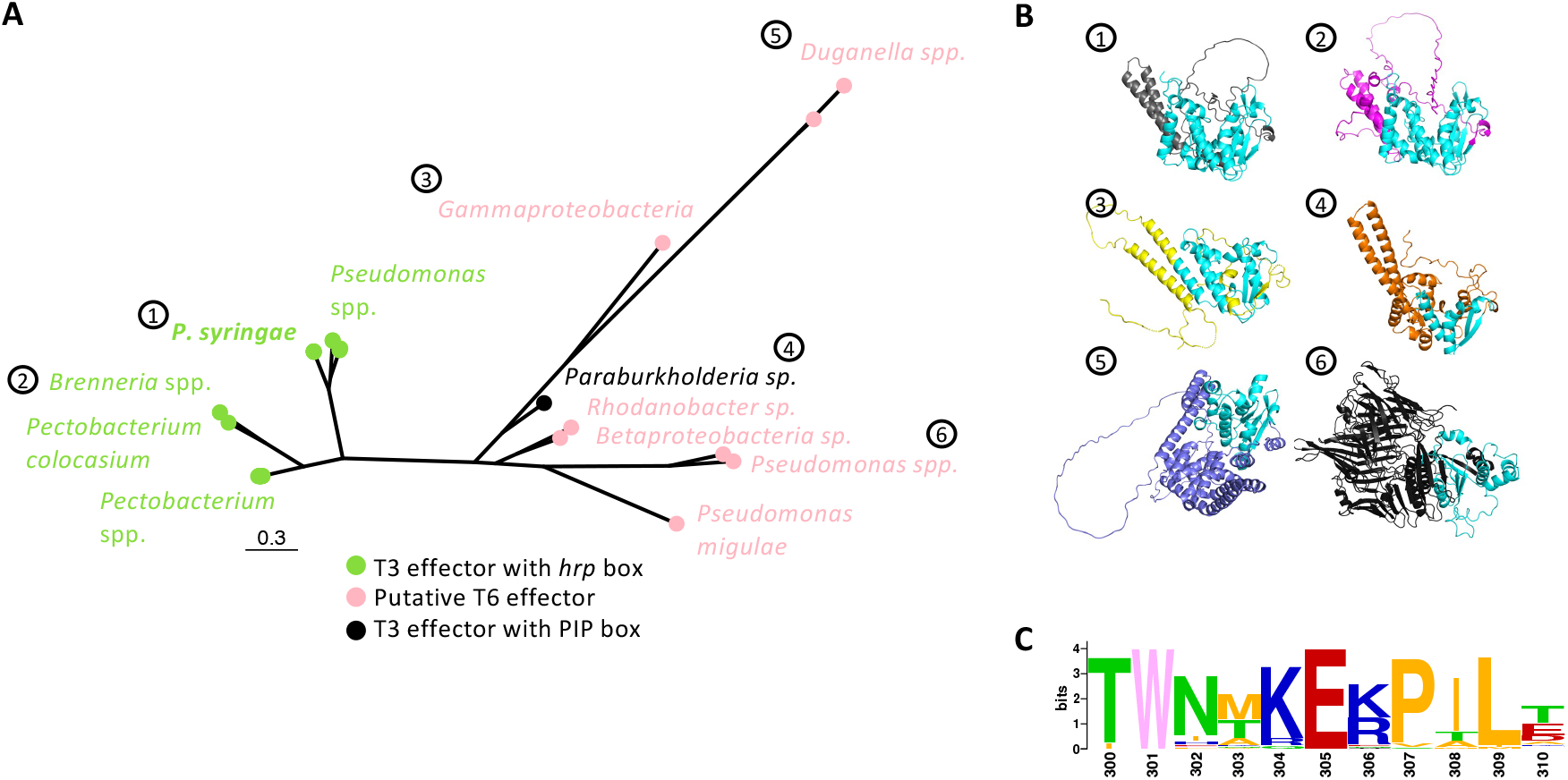
HopBY homologs are present in diverse bacterial species as type III or type VI effectors. A) Maximum-likelihood phylogeny of HopBY homologs using structure-guided protein sequence alignment. The homologs are grouped in two clades with the *P. syringae* HopBY labeled as “1”. Tips are labelled as green (T3Es with a *hrp* box in the promoter) or pink (putative T6Es). The predicted Hrp2 T3E was labelled as black. The scale shows substitutions per site. B) AlphaFold2 models of HopBY homologs, which show diverse overall structures. Numbers correspond to panel A. 1: *P. syringae coronafaciens* ICMP 4457, 2: *Brenneria* sp. EniD312, 3: *Gammaproteobacteria* RS470, 4: *Paraburkholderia* sp. ZP32-5, 5: *Duganella* sp. CF517, 6: *Pseudomonas frederiksbergensis* PgKB32. The HopBY-like ADPR cyclase domain is always on the C terminus of each protein, which is highlighted in cyan. **C)** Sequence logo showing the conservation of residues in the region surrounding the catalytic Glu residue (corresponding to positions 300-310 aa in the *P. syringae* HopBY). W301 and E305 are 100% conserved in all the 44 homologs.

T6S is an important secretion system for bacterial interactions with other bacteria and eukaryotes by delivering toxins that kill recipient cells (40). Indeed, these more distant homologs of the *P. syringae* HopBY were predicted to be putative T6S effectors. Some of these proteins have domains such as the N-terminal Rhs (Rearrangement Hotspot) core domain, which are often found in T6Es (41).

Using structure models predicted by AlphaFold2 (Fig 5B), we aligned the predicted ADPR cyclase domain in all the homologs to the *P. syringae* HopBY, which confirmed their structural similarity with RMSD values ranging from 0 - 4.44 Å (Table S5). We then aligned the region surrounding the catalytic residue E305 of *P. syringae* HopBY. This revealed that the glutamate as well as an upstream tryptophan, which was shown to be important for cyclization of the NAD^+^ hydrolysis products (6, 32), are highly conserved in all homologs, indicating they may be active enzymes (Fig 5C). The existence of HopBY ADPR cyclase domain homologs across effectors from T3 and T6 secretion systems indicates this enzymatic activity may have been adopted by distinct secretion systems that function in interspecies interactions.

## Discussion

Exploiting the pangenomic diversity and the extensive genome sequence resources of *P. syringae,* we conducted a comprehensive search to identify T3Es with enzymatic activities that can hydrolyse NAD^+^. This revealed host NAD^+^ manipulation as a virulence strategy in a model plant pathogen. *P. syringae* encodes a diverse set of T3Es with six different enzymatic activities, which potentially modulate NAD^+^ metabolism in plant cells during infection. Thus, manipulation of NAD^+^ and NAD^+^-related molecules is an important virulence mechanism commonly used by diverse *P. syringae* strains and likely by other plant and animal pathogens as well as bacteriophages.

NAD^+^ is a key metabolite that is involved in a wide range of cellular processes. NAD^+^ depletion could lead to cell death (4). Therefore, it is possible that effectors contribute to tissue damage through NAD^+^ depletion, which may be related to the development of disease symptoms. Indeed, the new T3E HopBY identified in this study caused chlorosis and tissue collapse. HopBY delivery by D36E led to a moderate, but statistically significant reduction in NAD^+^ abundance in infected tissue (Fig S8). Since not all plant cells in the leaf will receive effector delivery during infection, the observed reduction in overall NAD^+^ level may indicate significant depletion in those cells that do receive HopBY. Conceivably, HopBY-associated necrotic symptoms is due to NAD^+^ depletion in these cells. HopBY-associated phenotypes are EDS1-independent, indicating that they are not due to activation of EDS1-mediated plant immunity but may be a result of NAD^+^ depletion. Overexpression of the TIR domain of AbTIR and some plant TIRs in *N. benthamiana* also cause cell death independent of EDS1 (9, 42). This suggests alternative immune signalling pathways may exist, or that these TIR proteins may cause NAD^+^ depletion when accumulated to a high level.

In addition to the potential impact of NAD^+^ consumption activities, NADase effectors could also promote disease through the production of specific metabolites. The newly emerged role of NAD-derived small molecules in immune signalling leads to a possibility in which pathogens can use NADase effectors to suppress immunity. We can speculate that pathogen effectors could compete with host immune proteins for substrates (43). In addition to NAD^+^, the enzymatic domains identified from the *P. syringae* T3Es may also use other molecules such as ADPR and cADPR as substrates. Therefore, hydrolysis and depletion of immune signal molecules or their precursors could be a virulence mechanism. Although these hypotheses remain to be tested, investigation of the enzymatic activity of NADase effectors will offer a unique opportunity to dissect immune signalling through NAD-derived small molecules.

TIR proteins across biological kingdoms generate cyclic nucleotide products that play vital roles in immune signalling (3, 4). *P. syringae* produces HopAM1 as a TIR effector and HopBY shows structural and functional similarity to both TIR and ADPR cyclase. Both HopAM1 and HopBY could promote bacterial colonization indicating that they could either suppress plant immunity (15) and/or promote an environment suitable for disease progression. It is intriguing that plant and pathogen NADase activities can result in opposite phenotypes (resistance vs susceptibility). An attractive possibility is that these TIR/ADPR cyclase effectors produce distinct isomers that can activate immune signalling. Indeed, HopAM1 produces 3’cADPR, distinct from the 2’cADPR detected from the NAD^+^ hydrolysis reaction of activated plant TIR proteins (3) and could function as a negative regulator of plant immunity (6, 15). However, the somewhat surprising finding that HopBY produced a significant amount of 2’cADPR during bacterial infection reflects the complexity of NAD^+^ metabolism in immune response.

A role for 2’cADPR in TIR-NLR initiated immune signalling has been hypothesized but not formally demonstrated (3). Several previous observations suggest that 2’cADPR production alone is insufficient to activate immune signalling. A fusion receptor with the bacterial AbTIR and a mammalian NLR did not activate cell death when expressed in *N. benthamiana,* whilst several plant TIR-mammalian NLR fusions did (36). Furthermore, expression of bacterial TIRs alone led to 2’cADPR production but did not activate EDS1-mediated immunity in plants (4, 9). Finally, recent *in vitro* studies show that EDS1-helper NLR complexes directly bind to pRib-AMP, pRib-ADP, ADPr-ATP and diADPR. Therefore, although 2’cADPR can be produced by plant TIRs, it appears insufficient to directly initiate EDS1-mediated immune signalling.

Homologous proteins to HopBY were identified across diverse bacterial lineages. The *P. syringae* allele is closely related to other predicted T3Es from plant pathogens, but the other homologs were predicted as T6Es, which are involved in bacteria-bacteria and bacteria-eukaryotic interactions (40). NADase T6Es have been discovered before. Two ART-like families of T6Es, Tse6 and Tne2, were involved in bacteria-bacteria competition. Injecting into the recipient cells, Tse6 and Tne2 deplete NAD^+^, leading to cell death (45, 46). This mechanism is reminiscent to bacterial anti-viral immunity which involves TIR signalling to activate the sirtuin ThsA to deplete NAD^+^. It is intriguing that HopBY homologs are widespread in bacteria as T6Es. Testing whether these T6Es can function as cytotoxins in bacteria-bacteria or bacteria-eukaryote competitions will be an interesting future experiment. Presumably, the bacterial strains possessing T6 effectors related to HopBY must also encode an immunity protein or mechanism to prevent the action of the T6 effector before secretion (46). Some of the HopBY homologous T6Es contain the N-terminal Rhs domain (47). Rhs allows diversification of T6Es at the C-terminal toxin domains via recombination (41). The HopBY NADase domain likely recombined into these Rhs loci, generating a new virulence function. Similarly, the HopBY homologous T3Es could also be created through fusion to the T3S-dependent promoter and N-terminal secretion sequence via the terminal reassortment mechanism (48). Therefore, the HopBY domain has been incorporated into effectors for different interspecies competitions using distinct secretion system machinery, demonstrating the importance of NAD^+^ manipulation in these antagonistic interactions.

Much remains to be discovered about the diverse NADase activities of TIR domains and the complexity of their cyclic nucleotide products. The prevalence of pathogen effectors with NADase activities testifies to the importance of NAD^+^ metabolism as a battleground in host-pathogen interactions. Understanding how effectors with NAD^+^ hydrolysis activities promote virulence, and if and how they can disrupt immune signalling pathways in plant hosts, will continue to provide new insights into immune signalling and its circumvention in bacteria, animals, and plants.

## Methods

Standard procedures used in this manuscript are documented in detail in the Supplementary Methods. Specific methods developed here are included below.

### NADase T3E prediction

The pipeline to identify nucleosidase candidates is presented in Fig S2A. For HMMER (http://hmmer.org/) analysis, Pfam seed alignments for enzyme families (listed in Table S1) were obtained from Pfam (http://pfam.xfam.org/) (downloaded on December 1^st^, 2020). The script hmmbuild was used to create a HMM profile for each seed alignment. The command hmmsearch was used to screen all *P. syringae* proteins. HMMER results were filtered based on a tolerant bit-score of ≥ 15 (49) and grouped by orthogroup. Orthogroups with positive hits were further filtered to remove those in which less than 5% of the orthogroup members were a hit as these rare hits may not represent the orthogroup function.

HHsearch (50) was also used for each orthogroup to predict similarity to proteins in the PDB database (downloaded on October 28^th^, 2021). First, a multi-sequence alignment (MSA) was generated using ClustalW and a consensus sequence for each orthogroup was generated using hhcon. Next, the secondary structure predicted by PSIPRED was added to the alignment (51) before running HHsearch. Bash scripting was used to extract candidates with hits towards NADase enzymes in the database using a cutoff of ≥ 80% probability score. Two additional hits per protein with non-overlapping regions were also identified.

Candidate orthogroups predicted with HMMER or HHsearch were next screened with EffectiveT3 (29). Orthogroups containing hits with a score ≥ 0.9 were further analyzed for putative *hrp* box sequences in their promoter region (within 1000 bp upstream of the protein-coding sequence) using FIMO (30) based on the *hrp* box sequence GGAACC(15-17N)CCACNNA as input.

Candidate T3Es were manually filtered based on the percentage of sequences within the orthogroup predicted to be an effector (≥ 50%) and the percentage of these sequences that contained a possible *hrp*-box promoter (≥ 25%). Orthogroups only present in genomes without a predicted Hrp1 T3S were removed. Candidates T3Es were then manually examined to determine if they had characteristic N-terminal characteristics of *P. syringae* T3Es (31, 52). This included 1) a high percentage of serines in the first 50 amino acids; 2) an aliphatic amino acid in positions 3 or 4; and 3) a lack of negatively charged residues within the first 12 positions. Genomic locations of each T3E candidate gene were also manually examined using Geneious Prime 2022.0.1 to examine gene neighbourhoods (https://www.geneious.com).

### HPLC-MS

Cyclic-ADP ribose was analysed using a 1290 Infinity UHPLC system equipped with a 6546 Q-ToF (Agilent). Separation was on a 100×2.1mm 2.6μ Kinetex EVO C18 column connected in series to a 100×2.1mm 2.6μ Kintex F5 column (both Phenomenex) using the following gradient of methanol (solvent B) versus 0.1% formic acid adjusted to pH 6.0 with ammonium hydroxide (solvent A), run at 0.27 mL.min-1 and 35°C: 0 min, 0% B; 1.5 min, 3% B; 4.5 min, 15% B; 6.0 min, 60% B; 6.5 min, 95% B; 7.5 min, 95% B; 7.6 min, 0% B; 11.6 min, 0% B. Detection was by positive electrospray ionisation using the dual jet-stream ESI source. The instrument collected full MS from m/z 100-1700 at 6 spectra per second, and automatic MS/MS of the two most abundant precursors at 8 spectra per second, with an isolation width of m/z 4.0 (“medium”) and 35% collision energy. Each precursor was ignored for 0.2 min, in favour of less abundant precursors, after two MS/MS spectra had been collected.

Spray chamber conditions were 8 L.min-1 drying gas at 320°C, 35 psi nebulizer pressure, 11 L.min-1 sheath gas at 350°C, 3500 Vcap, 1000V nozzle voltage, 175V fragmentor, 65V skimmer, and 750V Octopole RF. The instrument was calibrated according to the manufacturer’s instructions before use, and reference masses (121.0509; 922.0098) were used during the run to ensure mass accuracy.

## Acknowledgments

We thank Dr. Alan Collmer for providing the *P. syringae* D36E mutant and Ma lab members for discussions and technical support. We thank Dan MacLean, Clara Jégousse and George Deeks for bioinformatic analysis support. Drs He Zhao and Hee-Kyung Ahn provided AbTIR expressing *E. coli eds1-2 Arabidopsis* seed and the pBBR1*MCS-5::avrRps4* plasmid.

## Funding

W.M. and J.J. are supported by Gatsby Charitable Foundation and UKRI BBSRC Grant BBS/E/J/000PR9797

## Competing Interests

The authors declare no competing interests.

## Data and material availability

All code are available at https://github.com/michhulin/Pseudomonas. Materials are available from W.M. upon request under a materials transfer agreement with the Sainsbury Laboratory.

## Notes

### Competing Interest Statement

The authors have declared no competing interest.

### Summary of Updates

This version of the manuscript includes further datasets regarding metabolomic analysis, functional characterisation and evolutionary analysis of effector protein HopBY

