## Supplementary materials for "Pangenomic analysis reveals plant NAD^+^ manipulation as an important virulence activity of bacterial pathogen effectors"

**This includes:**

- Supplemental methods
- 8 supplementary tables (1,2,6,7,8 in this file and 3,4 and 5 as excel)
- 10 supplementary figures
- References for the supplementary methods

#### Supplementary methods

##### Bacterial strains and plant material

Strains and plasmids used in this study are listed in Table S6. *Pseudomonas* strains used in this study include *P. syringae* pv. tomato strain DC3000 (PstDC3000), the T3S mutant Δ*hrcC* (1), and the effector-less mutant D36E (2). *P. coronafaciens* NCPPB 2397 (equivalent to ICMP 4457, genome assembly GCA_001400695) was used to clone the novel T3E OG18056. Antibiotics were used at the following concentrations kanamycin: 50 µg/mL, rifampicin: 50 µg/mL, carbenicillin: 100 µg/mL, gentamycin 50 µg/mL. *Agrobacterium tumefaciens* strain GV3101 was used for *Agrobacterium*-mediated transient expression in *N. benthamiana*.

*N. benthamiana* plants were grown in a controlled growth room with 16:8 light:dark conditions at 22 °C, 80% humidity. *A. thaliana* col-0 and *eds1-2* mutant plants were grown in a growth chamber with 10:14 light:dark conditions at 20 °C, 80% humidity. Five week old *N. benthamiana* and four week old *A. thaliana* were used for agro-infiltration and *P. syringae* inoculations.

##### Plasmid construction

Golden gate cloning (3) was used to generate constructs for transient expression of effectors in plants. Level 0 parts included: 35S long promoter, CDSNS (no stop) acceptor, 3xFLAG C-terminal tag, 35S terminator. The CDS were ordered from Twist Bioscience and PCR amplified using Phusion HF polymerase (ThermoFisher) to add golden gate compatible overhangs. Primers used in this study are presented in Table S7. CDS were inserted into the level 0 CDSNS acceptor using one-pot digestion-ligation reaction with the restriction enzyme BpiI-HF (New England Biolabs). Level 0 fragments were then assembled into the *in planta* expression vector pICH47732.

The *OG18056* effector gene was cloned into pUCP20TK for expression in *P. syringae*. Genomic DNA containing CDS and its native promoter (250 bp upstream of the start codon) was amplified from *P. coronafaciens* NCPPB 2397 with the addition of a C-terminal HA tag in the reverse primer. Infusion cloning was used to insert promoter and CDS fragments in reverse orientation to the *lacZ* promoter in the pUCP20TK vector after linearisation of the vector with BamH1 and EcoR1 (New England Biolabs).

For generation of the E305A mutant of OG18056, a site-directed mutagenesis approach using infusion was performed, splitting the gene and promoter fragment into two PCR products for cloning with the mutation introduced in primers R1 and F2. All constructs were confirmed by sanger sequencing. The recombinant plasmids were electroporated into *P. syringae* as described in (4) apart from recovery growth was for 2 hours at 28 °C, 220 rpm. The vectors pBBR1MCS-5 and pBBR1MCS-5::avrRps4 (5) were also introduced in *P. syringae* D36E via electroporation.

**Agro-infiltration in *N. benthamiana***

*A. tumefaciens* GV3101 with various constructs was streaked from glycerol stocks onto LBA plates containing appropriate antibiotics. Two-day old bacterial lawns were resuspended in sterile distilled water and washed twice by spinning at 3000 rpm for 10 minutes. The cells were resuspended in $MgCl_2$-2-(N-morpholino)ethanesulfonic acid (MES) buffer (10 mM $MgCl_2$, 10 mM MES, pH 5.6) to a final concentration of $OD_{600}$=1.0 before adding acetosyringone to a final concentration of 150 μM. Bacteria were incubated at room temperature (22 °C) for 3 hours before combining in a 1:1 ratio with the strain carrying the viral RNA silencing suppressor P19, meaning the final $OD_{600}$ of each *Agrobacterium* strain was 0.5. Protein expression was confirmed by immunoblotting using total proteins extracted from 1 cm leaf discs at 2 days post inoculation. Leaf discs were immediately frozen in liquid nitrogen and stored at -80 °C. To extract proteins, 100 μL of protein extraction buffer (50 mM Tris-HCl (pH 7.5), 150 mM NaCl, 1 mM EDTA, 0.1% Triton X-100) was added and the discs were ground with a micropestle whilst on ice. Crude plant material was removed by centrifugation at 11,000 x g for 15 minutes and 60 μL of the supernatant was combined with 30 μL 3X Laemmli buffer (6) with 1x cOmplete EDTA-free protease inhibitor cocktail (ThermoFisher) and boiled at 95 °C for 5 minutes. Immunoblotting was performed as described previously (7). An anti-FLAG antibody conjugated to horseradish peroxidase (Sigma) was used at 1:5000.

Immunoprecipitation of proteins was performed using 0.5 g of leaf material ground in liquid nitrogen and then 1 mL of protein extraction buffer (as above) added before centrifugation at 11,000 x g for 15 minutes. Input samples before addition of beads were included for comparison. 10 μL/mL of anti-FLAG M2 affinity Gel (A2220 Sigma-

Aldrich) was added to cleared samples and incubated at 4°C with gentle agitation for 2 hours. Beads were washed four times for 5 minutes in protein extraction buffer before resuspension in 60 µL protein extraction buffer. Immuno-precipitated proteins were visualised by immunoblot as above by boiling 20 µL of beads in Laemmli buffer.

**Effector expression and secretion analysis**

To confirm that novel effector protein OG18056 was induced in Hrp-inducing minimal medium and secreted in a T3S-dependent manner, PstDC3000 and the Δ*hrcC* mutant carrying pUCP20TK::OG18056-HA were used for protein extraction. A protocol based on (8) was utilised to extract secreted proteins from supernatants. The protocol was modified to extract from 8 mL of bacterial supernatant (rather than 2 mL). Immunoblotting was performed as in (7) using anti-HA from rat (Roche) at 1:1000, followed by a secondary anti-rat antibody conjugated to horseradish peroxidase (sc-2750, Santa Cruz) at 1:2000. The same protocol without supernatant extractions was used to confirm the expression of OG18056 and HopAM1 in PstDC3000 D36E used in bacterial infection assays.

***In planta Pseudomonas* pathogenicity assays**

PstDC3000 D36E carrying specific effector alleles or the pUCP20TK empty vector were grown from glycerol stock for 2 days on agar plates at 28 °C, and the lawn was resuspended in 10 mM $MgCl_2$ to the required optical density for inoculation. 1 x $10^8$ cfu/mL (OD 0.2) or 5 x $10^6$ cfu/mL (OD 0.01) was used to inoculate *A. thaliana* and monitor symptom development over several days, with pictures taken 1 or 3 days post inoculation. For bacterial population assays, inoculum was prepared at 5 x $10^5$ cfu/mL (OD 0.001) for *A. thaliana*. Bacteria were enumerated after 0 and 3 days by plating a dilution series on Kings B agar supplemented with rifampicin and kanamycin/gentamycin. All experiments were performed three times.

**NADase assay**

Semi-*in vitro* NADase assays were performed as in (9) with some modifications. After immuno-precipitation of proteins, 20 µL protein-laden FLAG beads were incubated with 6 µL 100 µM $NAD^+$ (Merck), 3.84 µL 10x PBS, 1 µL 0.6M NaCl, 1 µL 0.6M $MgSO_4$ and 28.16 µL water (total of 60 µL). The reactions were terminated after 1 hour at 28°C with shaking at 1000 rpm by pulling the beads to one side with a magnet and the supernatant was used directly for LCMS analysis. This experiment was repeated three times with similar results. The AbTIR 2'cADPR standard was extracted from induced *E. coli* lysate as in (10).

**Metabolite extraction *Arabidopsis thaliana* and LCMS analysis**

*P. syringae* D36E expressing effectors or just carrying the empty vector were inoculated into *A. thaliana* Col-0 leaves at OD 0.5 (~ 2.5 x $10^8$ CFU/mL) and harvested 18 hpi. Each sample consisted of 8 leaves harvested from two plants. Samples were freeze-dried for 48 hours. 0.01g of freeze-dried material was then extracted using 10% methanol 0.1% acetic acid as in (11). This experiment was repeated three times with similar results. $NAD^+$ depletion assays were performed as above but instead taking samples at day 0 and day 3 and the bacteria were inoculated at a lower dose (~ 5 x $10^6$ CFU/mL).

To visualise LCMS peaks Agilent software MassHunter Qualitative was utilised. The EIC of 542.06-542.07 m/z was extracted and data plotted for cADPR. For $NAD^+$ the EIC of 664.11-664.12 m/z was extracted. For quantitative analysis of $NAD^+$ the MassHunter Quantification software TOF (Quant-My-Way) was used. $NAD^+$ was set as the compound with a m/z of 664.11 with retention time of 4.2 minutes with allowance of 0.1 minutes either side. A calibration curve was generated with $NAD^+$ dilution series from 0.078 μM, 0.3125 μM, 1.25 μM, 5 μM, 10 μM, 15 μM, 20 μM $NAD^+$. Peaks were gaussian smoothed and areas were integrated automatically and then manually inspected and adjusted if necessary.

**Genome retrieval and annotation**

2,437 *P. syringae* (taxid136849) genomes were download from NCBI on November 18th, 2020. Genomes were grouped into 806 clusters showing ≥ 99.95% ANI using FastANI v1.32 (12). One genome per cluster was chosen based on the assembly with the lowest number of contigs. Genomes were filtered using an approach similar to Levy*, et al.* (13), removing those with ≥ 1000 contigs and an N50 value lower than 40,000 bp computed with Quast (14). Finally, CheckM (15) was used to select genomes ≥ 95% complete and < 5% contamination, leaving 531 genomes. Using a cut-off of ≥ 80% ANI, all but two genomes are clustered into a single group. These two genomes (GCA_ 900582625 and GCA_900573885) were kept as out-groups.

Genome annotation was performed using PROKKA (16) and Orthofinder (17) was used to define orthogroups. To generate a core genome phylogeny, 409 single copy orthogroups present in all genomes were aligned using ClustalW (18) with ends trimmed by Gblocks (19). Sequences were then concatenated into a 68,325 bp alignment, converted to the phylip format and input into IQ-TREE2 (20) with parameters JTT+I+G and 1000 bootstraps.

**Identification of Type III secretion system (T3S) and known Type III effectors (T3Es)**

To identify genomes containing a canonical *hrp* T3S, a BLAST-based approach was taken similar to that described in Dillon et al. (2019). T3S proteins from PstDC3000, *P. syringae* pv. *syringae* B728A, and *P. viridiflava* LP23.1a (Table S8 lists accession IDs) were used in a tBLASTn to query each genome. A genome was determined to putatively encode a *hrp* T3S if it had a hit to every query with > 30% length and > 30% identity for at least one of the three T3S inputs. A similar approach was taken to determine which orthogroups corresponded to known T3E families in *P. syringae*. A set of T3E protein sequences from pseudomonas-syringae.org was used as input to query all orthogroups with BLASTp. Orthogroups containing proteins with hit identities of 100% and E-value < $1x10^{-10}$ were determined to be those corresponding to known T3Es.

**Identification of homologues of OG18056**

The sequence of OG18056 from *P. coronafaciens* was searched against the non-redundant protein database on NCBI in August 2022, using BLASTP. All hits were extracted and full-length proteins were aligned using the structural-aware alignment program PROMALS-3D (21). The alignment was then used to create a phylogenetic tree using IQ-TREE (20) with model WAG+I+G4. To compare genomic regions, all genomes with a hit on BLASTP were downloaded and the genomic region was identified manually using Geneious Prime (2022.0.1). Upstream hrp (GGAACC(15-17N)CCACNNA) or PIP (TTCGC(15N)TTCGC) promoter motifs were identified as well as neighbouring gene ontologies. To identify if each genome possessed particular type 3 secretion systems, SctV homologues from five distinct T3SS (Q5K5P1_ECOLX, HRCV_RALSO, INVA_SALTY, Q9I327_PSEAE, Y4YR_SINFN, Q887B4_PSESM) were searched against each genome using tBLASTn. Present T3SS's were determined by assessing the best hits to each SctV homologue. To assess T6 effectors, all OG18056 homologues were uploaded to the SecReT6 online search engine (22) which predicts T6 effectors. In addition, *hopBY* homologues that were within T6 machinery gene clusters on the genome were also considered as possible T6 effectors.

**Protein structure prediction and alignment**

Structural prediction was conducted using ColabFold AlphaFold2 batch with MMseqs2 (23). Structural alignments were performed with fr-tm-align (24). This included alignments with hSARM1 (structure 6O0R) residues 561-700 aa and hCD38 (structure 6VUA). Aligned residues, root-mean-square deviation (RMSD) scores, and alignment lengths were extracted using bash scripting.

**Molecular docking of NAD$^+$**

Autodock Vina v1.2.3 (25) was used for molecular docking. P2Rank (26) was first used to predict catalytic pockets. Output files were processed with python to extract the pocket containing the catalytic residues that aligned with hSARM1-TIR. Pocket coordinates were added to the Vina config file for the docking run. The NAD$^+$ ligand was downloaded in 3D format from https://zinc20.docking.org (27) and converted to PDBQT format using the Autodock Vina prepare_ligand script. The Radius of Gyration was calculated and multiplied by 2.9 using python to generate the appropriate sized grid for ligand binding (28). Autodock Vina was then ran with exhaustiveness of 32 to produce eight possible models.

**Figure creation**

PyMOL (https://pymol.org/2/) was used for visualisation of protein structures. WebLogo (http://weblogo.threeplusone.com/) was used to generate a sequence logo of residues at particular positions in OG18056 homologs. Biorender (BioRender.com) was used to create diagrams. All other figures were created using R software (29).

Specifically, the packages ape (30), phangorn (31), gggenes (32), ggplot2 (33), ggtree (34) and reshape (35).

#### Statistical analysis

Analysis of variance (ANOVA) was used to statistically compare population counts of *P. syringae* from *Arabidopsis*. Tukey-HSD significance groups were extracted using R packages emmeans (36) and the cld function within agricolae (37) .

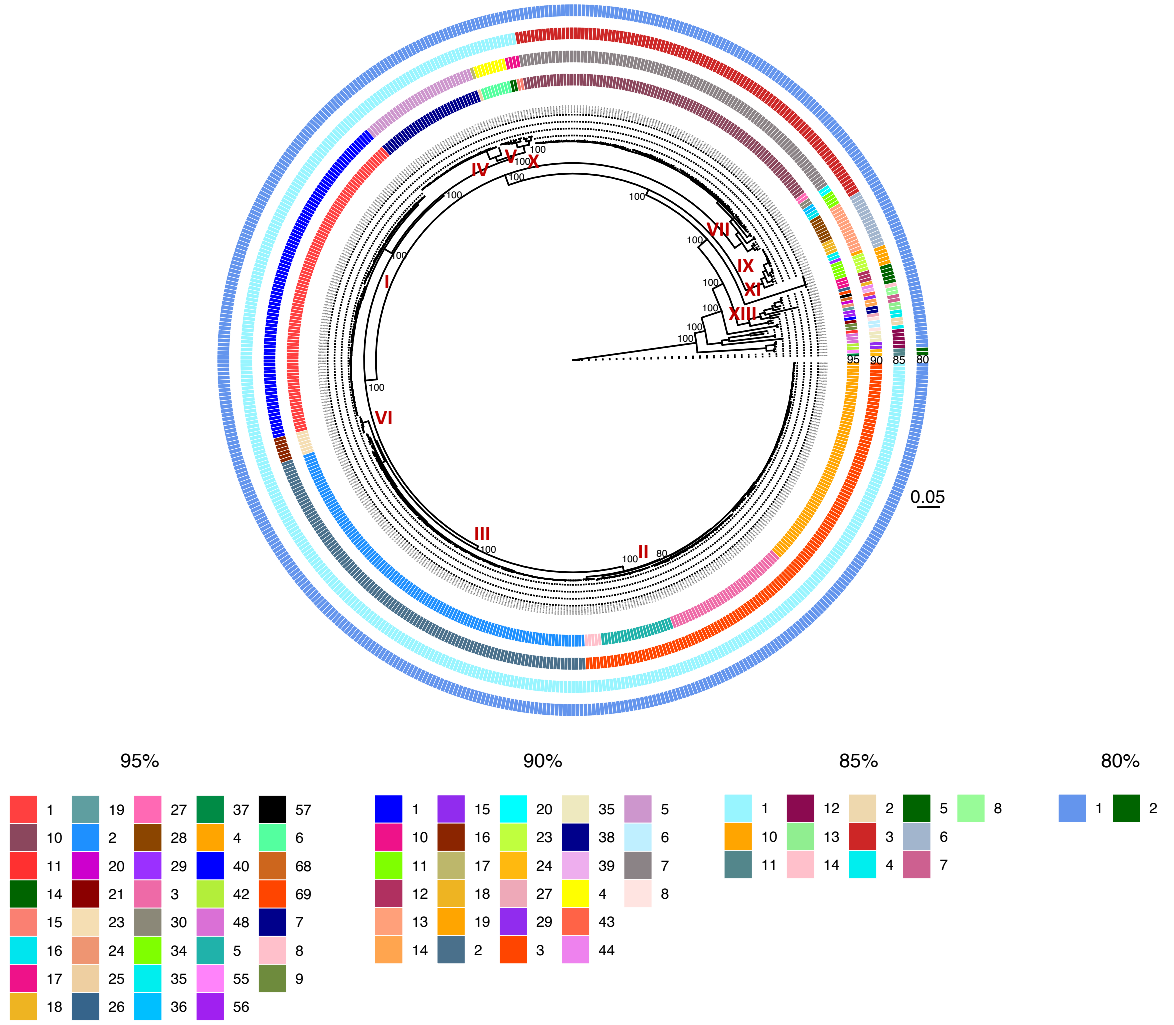


**Figure S1. Grouping *Pseudomonas syringae* by core genome and Average Nucleotide Identity (ANI).**

The maximum-likelihood core genome phylogeny was based on 409 single copy proteins encoded in all 531 genomes. Circles around the edge group genomes by Average Nucleotide Identity (ANI) calculated using FastANI at the 95%, 90%, 85% and 80% identity level. The corresponding groups are color-coded. Previously defined phylogroups are annotated onto the tree. The tree was rooted using *Pseudomonas* strains GCA_ 900582625 and GCA_900573885 that are divergent from *P. syringae*. The scale shows substitutions per site.

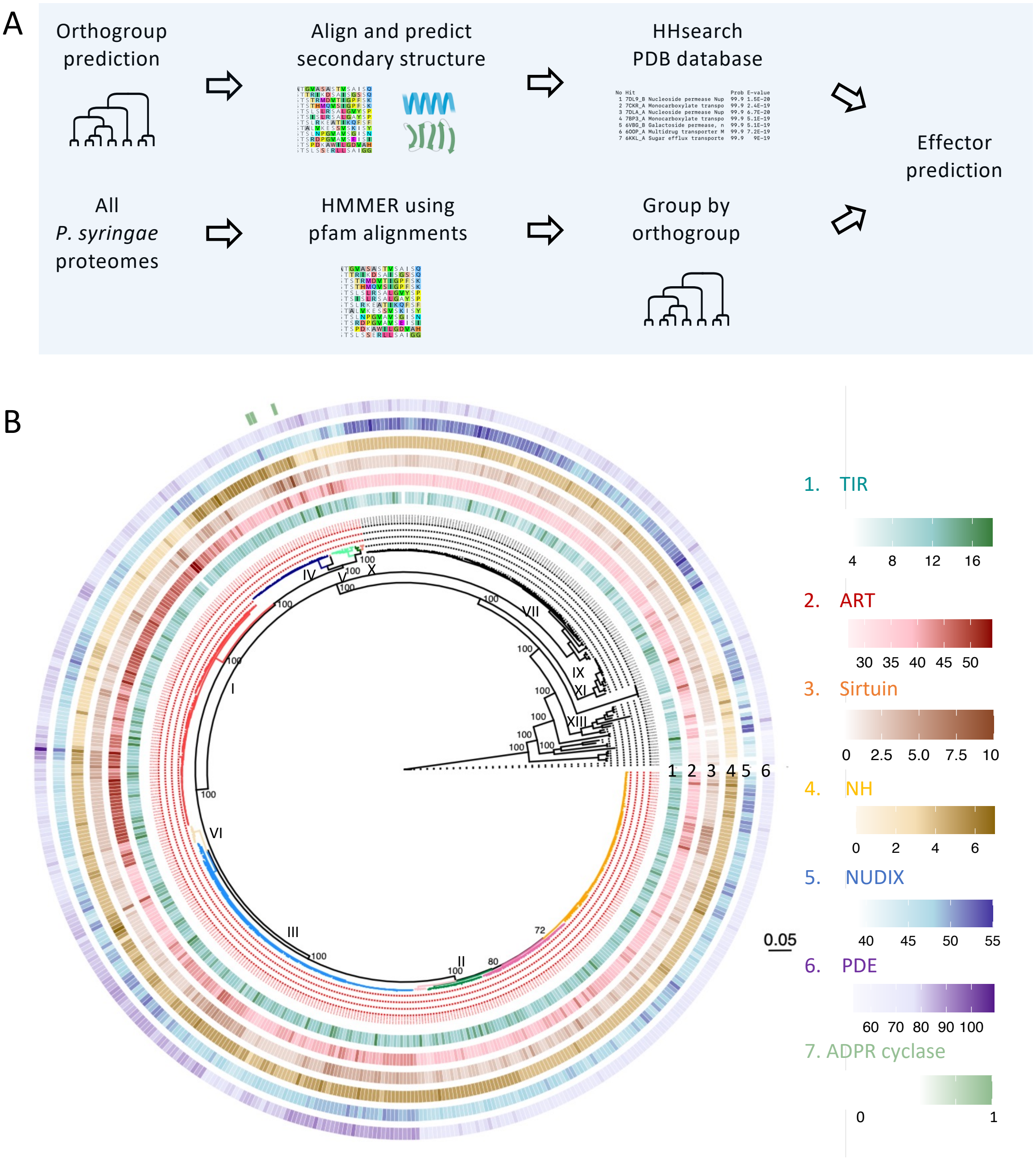


**Figure S2. Identification of NADases encoded in the *P. syringae* species complex.**

A. Bioinformatic pipeline illustrating the prediction of $NAD^+$ hydrolyzing enzyme families using HMMER and HHsearch in the *P. syringae* species complex.

B. Presence of NADases predicted across the *P. syringae* species complex is mapped onto the maximum-likelihood core genome phylogeny. Numbers of predicted proteins from each enzyme family is presented by heatmaps. Strains in the primary phylogroups, which mostly rely on the Hrp1 T3S for pathogenicity, are colored in red.

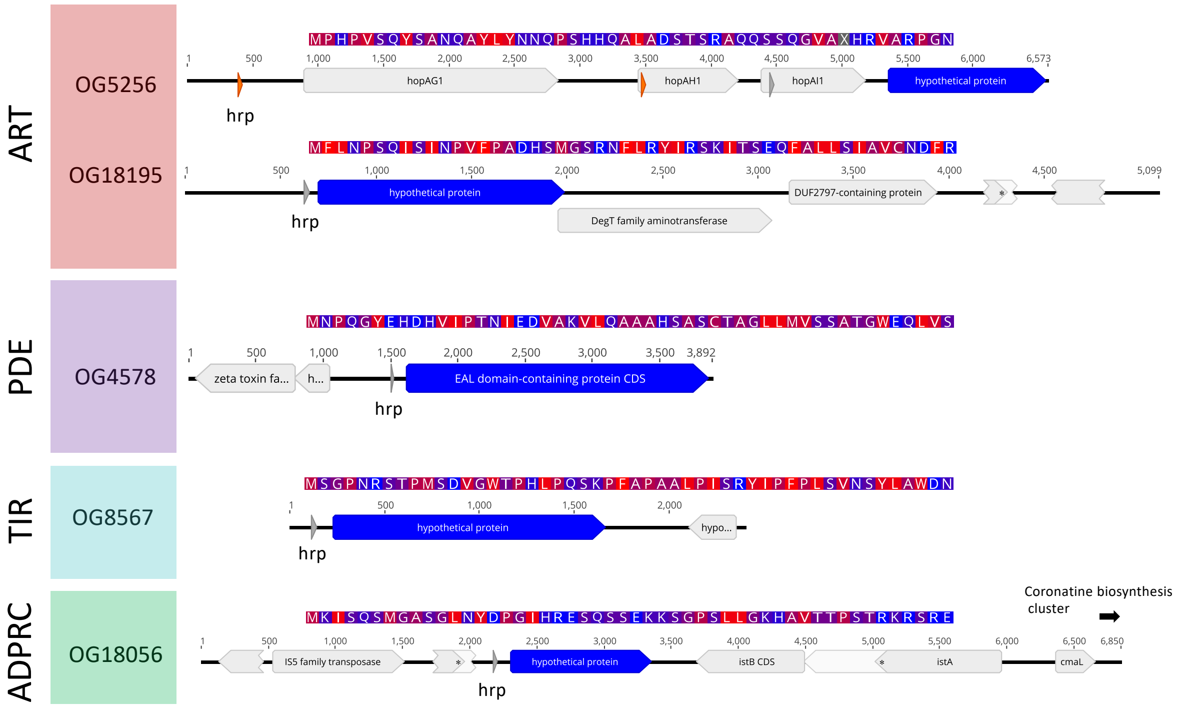


**Figure S3. N-terminal sequence predicted to be secretion signal (above) and the genomic location showing the predicted *hrp* box (below) of genes encoding the newly identified type III effector candidates.**

The type III-dependent secretion signal is predicted using the consensus amino acid sequence from a multiple sequence alignment of each effector candidate. The genomic location is from a representative genome. The location of *hrp* box in the promoter sequence is highlighted with a grey box. For OG5256, two possible alternative upstream *hrp* boxes (colored in orange) are shown as this gene may be part of an effector operon.

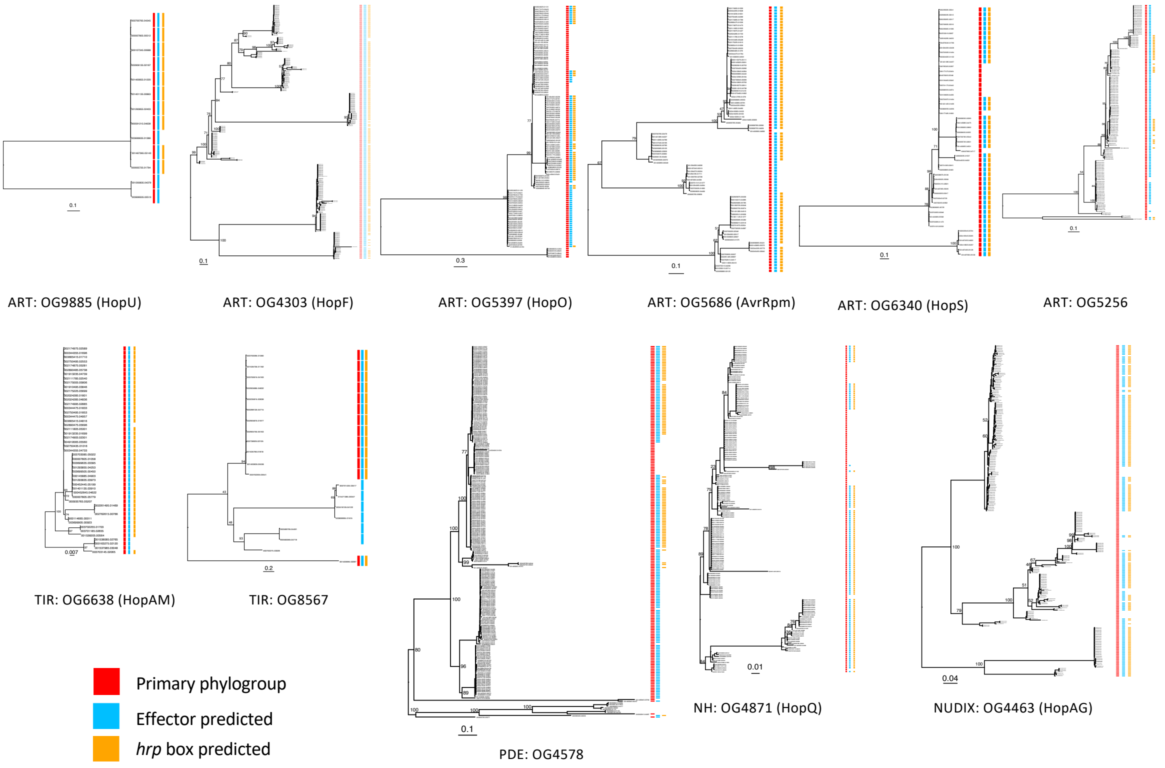


**Figure S4. Phylogenetic trees of NADase T3Es in *P. syringae*.**

Maximum-likelihood phylogenies were generated using IQ-TREE for all proteins in each orthogroup with < 50% gaps (to remove truncated proteins). The colored bars beside each tree label each sequence for: 1) whether it is from a primary phylogroup genome (red); 2) whether it was predicted as an effector (blue); and 3) whether it has a predicted *hrp* box in its promoter (orange).

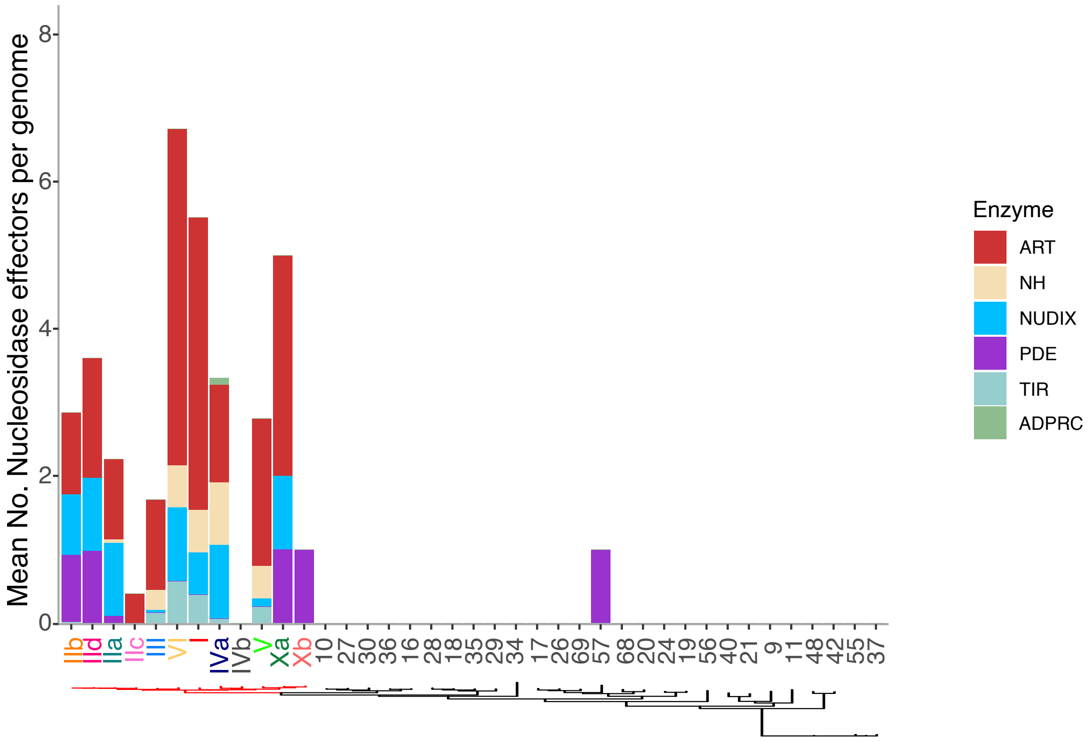


**Figure S5. Number of NADase T3Es encoded in each phylogroup of *P. syringae* species complex.**

The primary phylogroups are labelled with roman numerals. The Arabic numbers represent secondary phylogroups corresponding to their group number in Fig S1. The mean number of predicted NADase T3Es in each enzymatic family are shown in a collapsed version of the core genome phylogeny with the primary phylogroup clade branches labeled in red. ART: ADP-ribosyltransferase, NH: Nucleoside hydrolase, PDE: phosphodiesterase, ADPRC: ADP-ribose cyclase

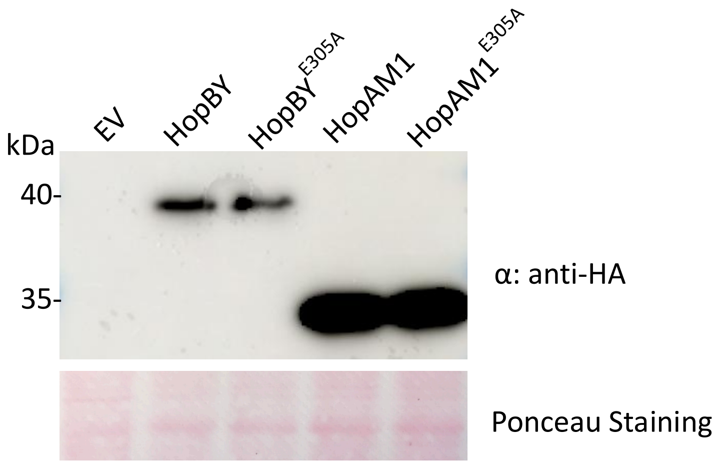


**Figure S6. Immunoblots showing the T3Es and their catalytic mutants were expressed in *P. syringae* mutant D36E.**

HopBY and HopAM1 were tagged with C-terminal HA. The genes were under their native promoters and cloned in the vector pUCP20TK. D36E carrying the recombinant plasmids were grown in Hrp-inducing minimal medium for 20 hours before cells were collected for western blotting using anti-HA antibody.

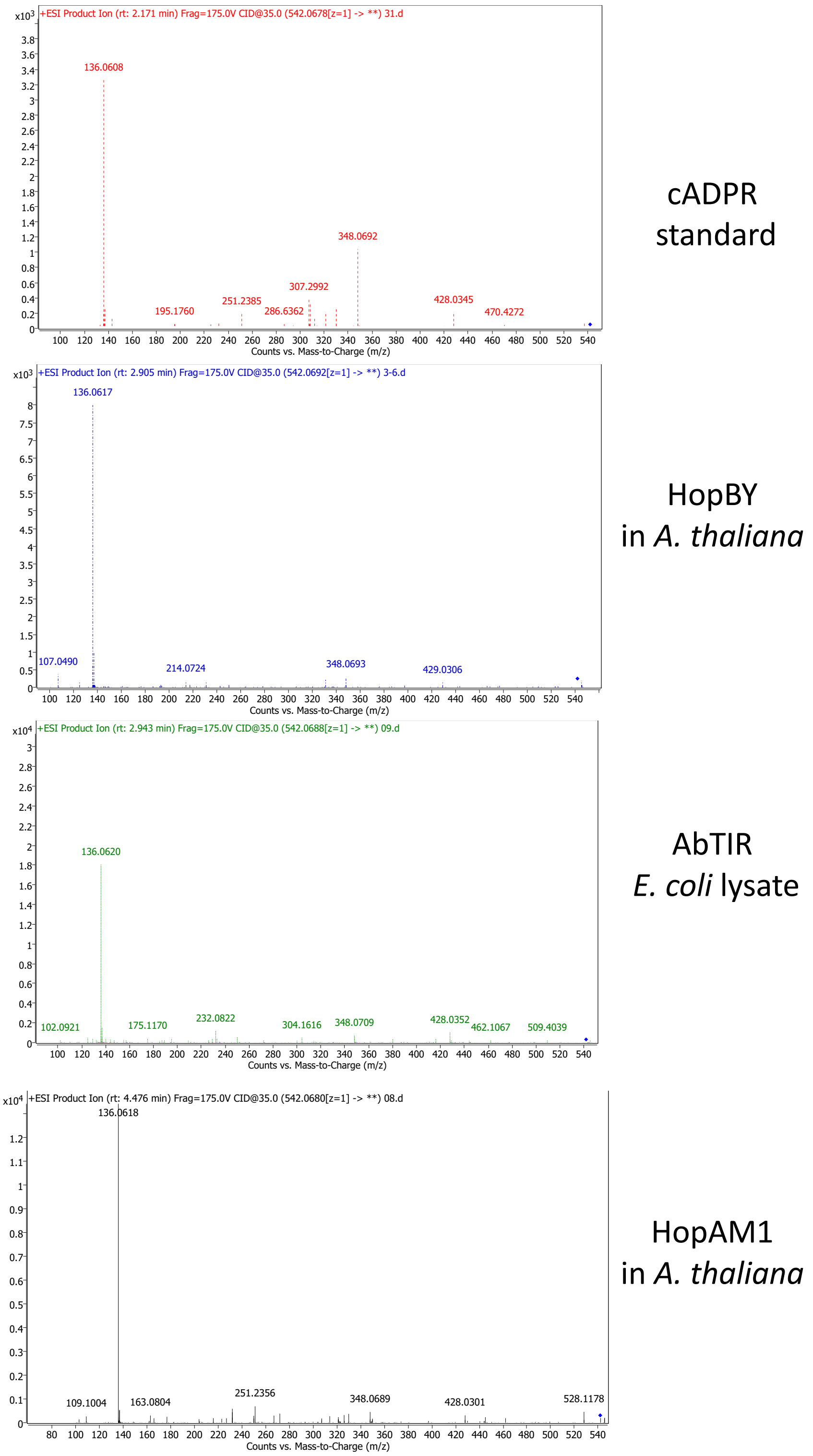


**Figure S7. MS/MS spectra of product ion scans of *m/z* 542.06-542.07.**

The samples include the cADPR standard, HopBY (*A. thaliana* leaf tissue inoculated with *P. syringae* D36E expressing HopBY), AbTIR (*E. coli* lysate) and HopAM1 (*A. thaliana* leaf tissue inoculated with *P. syringae* D36E expressing HopAM1). The blue diamonds in the spectra represent *m/z* 542.06.

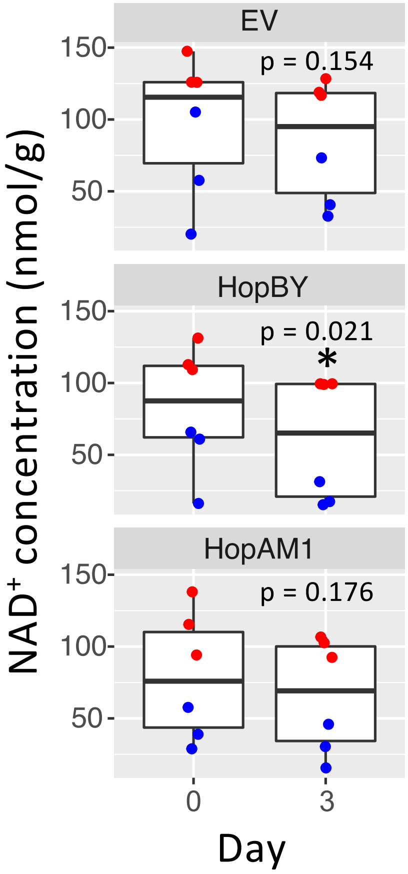


**Figure S8: NAD$^+$ quantification in *A. thaliana* leaves inoculated with D36E expressing HopBY, HopAM1 or carrying the empty vector at day 0 and 3 post inoculation.** NAD$^+$ was quantified using LCMS using a concentration curve of NAD$^+$. * denotes statistical significance using a t-test ($p<0.05$). The p value for each treatment is presented. The analysis is based on results of two independent experiments with data points colored in red and blue respectively.

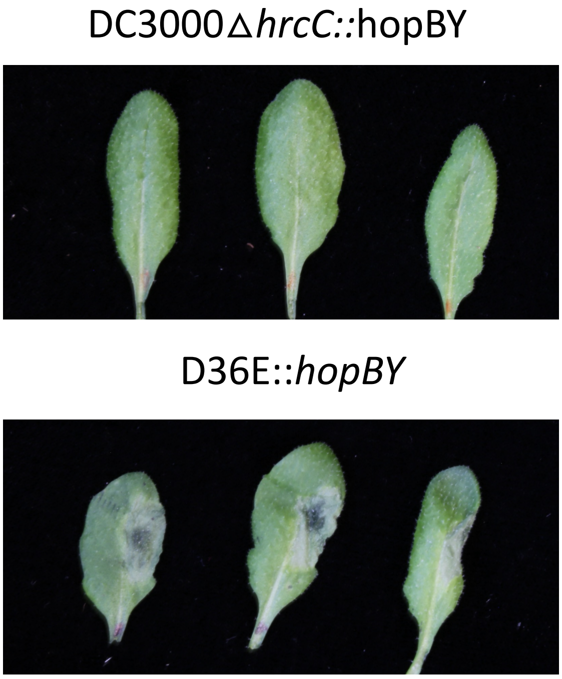


**Figure S9. HopBY symptoms are dependent on a functional T3 secretion system**
Representative images of *A. thaliana* leaves inoculated with either DC3000∆*hrcC::*hopBY or D36E::*hopBY* at a concentration of 1 x $10^8$ CFU/mL and imaged 24 hours post inoculation.

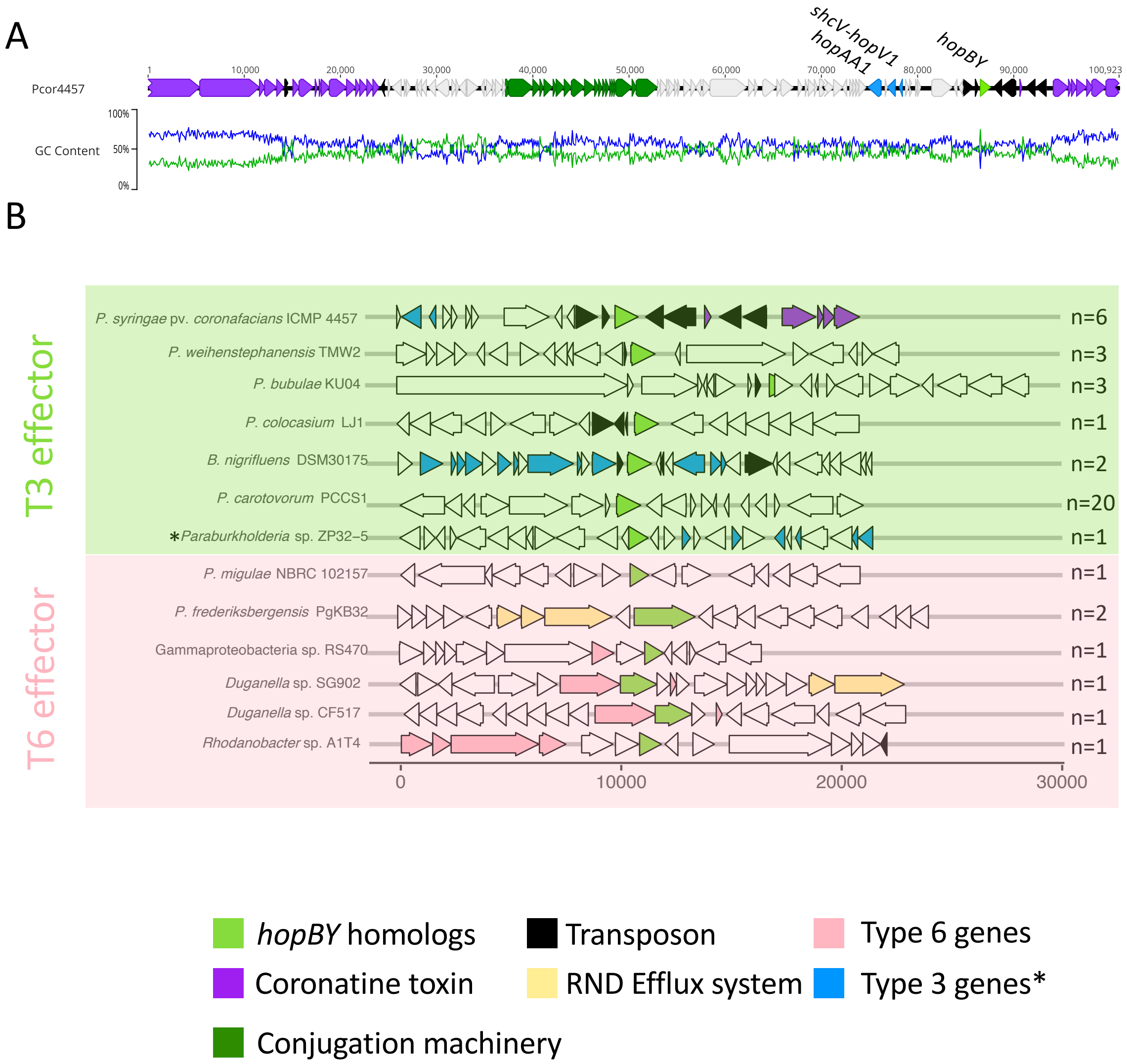


**Figure S10. Genomic location of the *hopBY* homologs.**

A. Genome region map of *hopBY* in ParICMP4457_Contig_64. Open reading frames are colour-coded to indicate functions. GC content is shown with GC (blue) and AT (green) percentage determined over a sliding window of 48 nucleotides.

B. Genome regions containing *hopBY* homologs in various bacteria. Gene content within a 30 Kb region surrounding the *hopBY* homolog (in green) is presented for each genome. The number of genomes containing the same gene layout is also shown. Open reading frames are color-coded to indicate functions. Genomes containing *hopBY* homologs as T3E or T6Es are labelled. The asterisk labels *Paraburkholderia* sp. Zp32-5, which encodes a *hopBY* homolog predicted to be a T3E although this bacterium has the Hrp2 T3S. All other bacteria that encode *hopBY* homologs as T3E have the Hrp1 T3S.

#### Table S1 PFAM accessions for each enzyme group used to construct HMM

| PFAM | Group | Description | Seed alignment size |
|---|---|---|---|
| **PF01582** | TIR | Toll-Interleukin receptor | 23 |
| **PF08937** | TIR | MTH538 TIR-like domain | 27 |
| **PF10137** | TIR | Predicted nucleotide-binding protein containing TIR-like domain | 49 |
| **PF13676** | TIR | TIR domain | 458 |
| **PF18567** | TIR | Toll-Interleukin receptor | 26 |
| **PF02267** | ADPR cyclase | Rib_hydrolayse | 27 |
| **PF01129** | ART | NAD(P)+-protein-arginine ADP-ribosyltransferase | 8 |
| **PF01375** | ART | Heat-labile enterotoxin family | 6 |
| **PF01885** | ART | RNA 2'-phosphotransferase, Tpt1 / KptA family | 222 |
| **PF02027** | ART | RolB/RolC glucosidase family | 13 |
| **PF00644** | ART | Poly(ADP-ribose) polymerase catalytic domain | 27 |
| **PF02917** | ART | Pertussis toxin | 2 |
| **PF03496** | ART | ADP-ribosyltransferase exoenzyme | 21 |
| **PF06108** | ART | Protein of unknown function (DUF952) | 188 |
| **PF08808** | ART | RES domain | 180 |
| **PF09009** | ART | Exotoxin A catalytic | 2 |
| **PF09143** | ART | AvrPphF-ORF-2 | 4 |
| **PF09475** | ART | Dot/Icm secretion system protein | 3 |
| **PF10386** | ART | Protein of unknown function (DUF2441) | 4 |
| **PF12120** | ART | Rifampin ADP-ribosyl transferase | 32 |
| **PF13151** | ART | Protein of unknown function (DUF3990) | 86 |
| **PF14021** | ART | Tuberculosis necrotizing toxin | 100 |
| **PF14487** | ART | ssDNA thymidine ADP-ribosyltransferase, DarT | 47 |
| **PF15633** | ART | HYD1 signature containing ADP-ribosyltransferase | 6 |
| **PF18648** | ART | Tse2 ADP-ribosyltransferase toxins | 5 |
| **PF18760** | ART | ADP-Ribosyltransferase in polyvalent proteins | 15 |
| **PF19048** | ART | SidE mono-ADP-ribosyltransferase domain | 3 |
| **PF00293** | NUDIX | NUDIX domain | 127 |
| **PF03559** | NUDIX | NDP-hexose 2,3-dehydratase | 70 |
| **PF09296** | NUDIX | NADH pyrophosphatase-like rudimentary NUDIX domain | 36 |
| **PF13869** | NUDIX | Nucleotide hydrolase | 17 |
| **PF14443** | NUDIX | DBC1 | 23 |
| **PF14815** | NUDIX | NUDIX domain | 384 |
| **PF15916** | NUDIX | Domain of unknown function (DUF4743) | 30 |
| **PF16705** | NUDIX | NUDIX, or N-terminal NPxY motif-rich, region of KRIT | 7 |

| | | | |
|---|---|---|---|
| **PF01156** | Nucleoside N-ribohydrolase | Inosine-uridine preferring nucleoside hydrolase | 664 |
| **PF02146** | Sirtuin | Sir2 family | 18 |
| **PF00149** | PDE | Calcineurin-like phosphoesterase | 324 |
| **PF00233** | PDE | 3'5'-cyclic nucleotide phosphodiesterase | 247 |
| **PF01663** | PDE | Type I phosphodiesterase / nucleotide pyrophosphatase | 67 |

#### Table S2 Number of orthogroups with NADase domains identified in *P. syringae*

| | HHsearch | HHsearch unique | HMM | HMM unique | Shared | Complete | Core | Accessory |
|---|---|---|---|---|---|---|---|---|
| TIR | 65 | 33 | 34 | 2 | 32 | **67** | 5 | 62 |
| NUDIX | 109 | 81 | 30 | 2 | 28 | **111** | 38 | 73 |
| ART | 140 | 112 | 34 | 6 | 28 | **146** | 28 | 118 |
| PDE | 226 | 186 | 40 | 0 | 40 | **226** | 53 | 173 |
| NH | 8 | 3 | 5 | 0 | 5 | **8** | 2 | 6 |
| SIR | 15 | 12 | 3 | 0 | 3 | **15** | 2 | 13 |
| ADPRC | 1 | 1 | 0 | 0 | 0 | **1** | 0 | 1 |
| Total | 564 | 428 | 146 | 10 | 136 | **574** | 128 | 446 |

#### Table S3. Metainformation for orthogroups identified as putative NADases in the *P. syringae* pangenome (see excel)

#### Table S4. NADase effectors identified using EffectiveT3 from the *P. syringae* pangenome (see excel)

#### Table S5. OG18056 homologues across bacteria are type III or type VI effectors (see excel)

#### Table S6 Bacterial strains and constructs used in this study

| Strain/construct | Purpose | Reference |
|---|---|---|
| *P. syringae* pv. tomato DC3000 | Expression, secretion assay, rifR | (38) |
| *Pst* DC3000 Δ*hrcC* | Expression, secretion assay, rifR | (1) |
| *Pst* DC3000 D36E | Virulence assays, rifR | (2) |
| *P. coronafaciens* NCPPB 2397 | Source of gDNA for cloning effector | (39) |
| GV3101 | Agroinfiltration, gmR, rifR | (40) |
| GV3101:P19 | *Agrobacterium tumefaciens* with silencing suppressor for agroinfiltration | (41) |
| DH10β/StellarTM competant cells | Cloning | ThermoFisher, Takara |
| pucp20TK | Broad host-range vector for *Pseudomonas* expression, kmR | (42) |
| pucp20TK:OG18056-HA | Expression of effector in *Pseudomonas*, kmR | This study |
| pucp20TK:OG18056-E305A-HA | Expression of effector in *Pseudomonas*, kmR | This study |
| pucp20TK:hopAM1-HA | Expression of effector in *Pseudomonas*, kmR | This study |
| pucp20TK:hopAM1-E191A-HA | Expression of effector in *Pseudomonas*, kmR | This study |
| pBBR1MCS-5 | Broad host range vector for Pseudomonas expression, gmR | (43) |
| pBBR1MCS-5: avrRps4 | Expression of effector in *Pseudomonas*, gmR | (5) |
| pICH51266 | 35s promoter, spR | (3) (addgene 50267) |
| pICSL01005 | Level 1 acceptor CDS no stop codon, carbR | (3) (addgene 174575) |
| pICSL50007 | 3xFlag (C-terminal), spR | (3) (addgene 50308) |
| pICH41414 | 35s terminator, spR | (3) (addgene 50337) |
| pICH47732 | Level 0 acceptor, spR | (3) (addgene 48000) |
| pICSL01005::35s-OG18056-3xFLAG | Transient expression of effector in plants, carbR | This study |
| pICSL01005::35s-OG18056-E305A-3xFLAG | Transient expression of effector in plants, carbR | This study |
| StrepTag Ab-TIR histag in pET30a+ | AbTIR expression in *E.coli* | (10) |

### Table S7 Primers used in this study

| Primer | Purpose | Experiment | Sequence (5'-3') | Amplicon size (bp) |
|---|---|---|---|---|
| hopAM1_cdsF | Add infusion overhangs to synthesised gene product with HA tag | Bacterial expression | GGAGTTCAAGATGCACGCAAATCCT | 888 |
| hopAM1_cdsR | | | TATGACCATGATTACGAATTCTAAG | |
| hopAM1_promF | Amplify promoter with infusion overhangs | Bacterial expression | CAGGTCGACTCTAGAGGATCCAGGCAAAGTAGATGCCGCGT | 246 |
| hopAM1_promR | | | TAAAGGATTTGCGTGCATGTATGCCTCCAGACGTT | |
| OG18056_F | Amplify promoter and CDS with infusion overhangs and add HA tag | Bacterial expression | CGACTCTAGAGGATCTTTGTCTGCACCCTCTTAGGAAAAA | 1359 |
| OG18056_HA_R | | | CCATGATTACGAATTCTAAGCGTAATCTGGAACATCGTATGGGT ACGTCAGGTTGCCACGAT | |
| OG18056_R1 E305A | Site-directed mutagenesis of OG18056 | Bacterial expression | GGATGGGCCGCGCCTTGGTGTTCCATGT | 1176 (with OG18056_F) |
| OG18056_F2 E305A | Site-directed mutagenesis of OG18056 | Bacterial expression | AGGCGCGGCCCATCCTCGACAGAAACAA | 185 (with OG18056_HA_R) |

**Table S8 Protein accession numbers for Hrp1 T3S identification**

| Strain | Accession | Description |
|---|---|---|
| *Pst* | AAO54900 | membrane-bound lytic murein transglycosylase D |
| *Pst* | AAO54901 | type III transcriptional regulator HrpR |
| *Pst* | AAO54902 | type III transcriptional regulator HrpS |
| *Pst* | AAO54903 | type III helper protein HrpA1 |
| *Pst* | AAO54904 | type III helper protein HrpZ1 |
| *Pst* | AAO54905 | type III secretion protein HrpB |
| *Pst* | AAO54906 | type III secretion protein HrcJ |
| *Pst* | AAO54907 | type III secretion protein HrpD |
| *Pst* | AAO54908 | type III secretion protein HrpE |
| *Pst* | AAO54909 | type III secretion protein HrpF |
| *Pst* | AAO54910 | type III secretion protein HrpG |
| *Pst* | AAO54911 | type III secretion protein HrpT |
| *Pst* | AAO54912 | negative regulator of hrp expression HrpV |
| *Pst* | AAO54913 | RNA polymerase sigma factor HrpL |
| *Pst* | AAO54926 | type III helper protein HrpK1 |
| *Pst* | AAO54927 | HrpK |
| *Pss* | AAY36238.1 | HrpL |
| *Pss* | AAY36244.1 | HrpJ |
| *Pss* | AAY36245.1 | HrcV |
| *Pss* | AAY36246.1 | HrcN |
| *Pss* | AAY36247.1 | HrpV |
| *Pss* | AAY36248.1 | HrpT |
| *Pss* | AAY36249.1 | HrpG |
| *Pss* | AAY36250.1 | HrpF |
| *Pss* | AAY36251.1 | HrpE |
| *Pss* | AAY36252.1 | HrpD |
| *Pss* | AAY36253.1 | HrpB |
| *Pss* | AAY36254.1 | HrpA |
| *Pss* | AAY36255.1 | HrpS |
| *Pss* | AAY36256.1 | HrpR |
| *Pss* | AAY36271.1 | membrane-bound lytic murein transglycosylase-like protein |
| *Pss* | AAY36272.1 | type III helper protein HrpW1 |
| *P.vir* | AAT96136.1 | type III transcriptional regulator HrpR |
| *P.vir* | AAT96137.1 | type III transcriptional regulator HrpS |
| *P.vir* | AAT96138.1 | type III helper protein HrpA2 |
| *P.vir* | AAT96139.1 | type III helper protein HrpZ1 |
| *P.vir* | AAT96141.1 | type III secretion protein HrpB |
| *P.vir* | AAT96150.1 | type III secretion apparatus lipoprotein, YscJ/HrcJ family |
| *P.vir* | AAT96151.1 | type III secretion protein HrpD |
| *P.vir* | AAT96153.1 | type III secretion protein HrpE |
| *P.vir* | AAT96154.1 | type III secretion protein HrpF |

| | | |
|---|---|---|
| *P.vir* | AAT96155.1 | type III secretion protein HrpG |
| *P.vir* | AAT96156.1 | outer-membrane type III secretion protein HrcC |
| *P.vir* | AAT96158.1 | type III secretion protein HrpT |
| *P.vir* | AAT96160.1 | negative regulator of hrp expression HrpV |
| *P.vir* | AAT96161.1 | Sigma-70 region 2:Sigma-70 region 4 |
| *P.vir* | AAT96162.1 | type III helper protein HrpK1 |
| *P.vir* | AAT96163.1 | outer-membrane type III secretion protein HrcC |
